# The Unique Evolutionary Trajectory and Dynamic Conformations of DR and IR/DR- coexisting Plastomes of the Early Vascular Plant Selaginellaceae (Lycophyte)

**DOI:** 10.1101/505867

**Authors:** Hong-Rui Zhang, Qiao-Ping Xiang, Xian-Chun Zhang

## Abstract

Both direct repeats (DR) and inverted repeats (IR) are documented in the published plastomes of four *Selaginella* species indicating the unusual and diverse plastome structure in the family Selaginellaceae. In this study, we newly sequenced complete plastomes of seven species from five main lineages of Selaginellaceae and also re-sequenced three species (*S. tamariscina, S. uncinata* and *S. moellendorffii*) to explore the evolutionary trajectory of Selaginellaceae plastomes. Our results showed that the plastomes of Selaginellaceae vary remarkably in size, gene contents, gene order and GC contents. Notably, both DR and IR structure existed in the plastomes of Selaginellaceae with DR structure being an early diverged character. The occurrence of DR structure was right after the Permian-Triassic (P-T) extinction (ca. 246 Ma) and remained in most subgenera of Selaginellaceae, whereas IR structure only reoccurred in the most derived subg. *Heterostachys* (ca. 23 Ma). The presence of a pair of large repeats *psb*K-*trn*Q, together with DR/IR region in *S. bisulcata, S. pennata, S. uncinata*, and *S. hainanensis*, could frequently mediate diverse homologous recombination and create approximately equal stoichiometric isomers (IR/DR-coexisting) and subgenomes. High proportion of repeats is presumably responsible for the dynamic IR/DR-coexisting plastomes, which possess a lower synonymous substitution rate (d*S*) compared with DR-possessing plastomes. We propose that the occurrence of DR structure, together with few repeats, is possibly selected to adapt to the environmental upheaval during the P-T crisis and the IR/DR-coexisting plastomes also reached an equilibrium in plastome organization through highly efficient homologous recombination to maintain stability.

**Data deposition:** All the plastomes were deposited in GenBank under accession numbers MG272483-MG272484, MH598531-MH598537 and MK156800.

## Introduction

Plastid genomes (plastomes) of almost all land plants are highly conserved and present the canonical quadripartite structure with a pair of large inverted repeats (termed IR_A_ and IR_B_) separated by two single copy regions (termed LSC and SSC) (Mower and Vickrey 2018). Normally, the range of the IR varies through expansion or contraction. Complete loss of the IR is rare but has been observed in some species of Fabaceae (Cai et al. 2008; Lavin et al. 1990), Geraniaceae (Blazier et al. 2016; Guisinger et al. 2011; Ruhlman et al. 2017), and Cactaceae (Sanderson et al. 2015). Remarkably, plastomes with a pair of large direct repeats (termed DR_A_ and DR_B_) have been documented for two species of Selaginellaceae, *Selaginella tamariscina* (Xu et al. 2018) and *S. vardei* (Unpublished data) in land plants. The DR structure in Selaginellaceae was explained to have occurred by ca. 50 kb fragment inversion with a complete IR_B_ being included, compared with the plastome of its sister family Isoetaceae (Unpublished data).

In addition to the exceptional existence of plastomes with DR structure, a salient fraction of land plants plastomes experienced significant structural rearrangements, with evidence of large inversions and loss of entire gene family, despite the overall conservation in structures and gene order (Mower and Vickrey 2018). A 30 kb inversion (*ycf*2-*psbM*) was detected in the large single copy (LSC) of ferns and seed plants plastomes relative to bryophytes and lycophytes, which is a strong evidence showing lycophytes are located at the basal position of vascular plants (Raubeson and Jansen 1992). The plastomes of ferns underwent two hypothetical inversions (CE inversion (*trnC* to *trnE*) and DE inversion (*trnD*-*trnY*)) within *rpoB*-*psbZ* (BZ) region from the ancestral gene order in eusporangiates to the derived gene order in core leptosporangiates whereas the plastome structures within these two groups were generally consistent, respectively (Gao et al. 2013; Gao et al. 2011; Grewe et al. 2013). Many other rearrangements also exist in some conifers (Chaw et al. 2018) and several angiosperm lineages like Campanulaceae, Fabaceae, and Geraniaceae (Mower and Vickrey 2018; Ruhlman and Jansen 2018). Inversion facilitated by recombination, transposition, and expansion/contraction of the IR have been suggested as three different mechanisms that cause rearrangements in land plants (Jansen and Ruhlman 2012). However, it is recently recognized that most plastomes exists as linear/concatemeric/highly branched complex molecules in plants and these rearrangement events are reinterpreted as result of a BIR-like, recombination-dependent replication mechanism between different linear plastome templates (Oldenburg and Bendich 2015). Furthermore, four families of nuclear-encoded proteins (MutS homologue 1 (*MSH*1), RecA-like recombinases, the organellar ssDNA-binding proteins (*OSB*s), and the Whirlies) have been characterized to target to both mitochondria and plastid, or some protein members of the four families target to only plastids and function as recombination surveillance machinery in plant plastids (Maréchal and Brisson 2010).

Gene and intron content are highly conserved among the vast majority of land plants plastomes, however, numerous examples of gene loss or pseudogenization have been identified in several angiosperm lineages (Ruhlman and Jansen 2014). For example, most or all of the suite of 11 functionally related *ndh* genes have been lost independently in a small assortment of taxa with diverse habitat, including the parasitic *Epifagus* (Wicke et al. 2011), the mycoheterotrophic *Rhizanthella gardneri* (Delannoy et al. 2011), some members of the carnivorous Lentibulariaceae (Wicke et al. 2013), the xerophytic *Suguaro cactus* (Sanderson et al. 2015) and gnetales (Braukmann et al. 2009), the aquatic *Najas flexilis* (Peredo et al. 2013) and some taxa with less unusual life histories, such as Pinaceae (Braukmann et al. 2009) and *Erodium* (Blazier et al. 2011a). In addition, the loss of protein coding genes and tRNA genes has occurred sporadically in different land plants lineages (Ruhlman and Jansen 2014; Wu and Chaw 2015). The fatty acid synthesis related gene, *accD*, has been lost from plastomes of angiosperm at least seven times (Jansen et al. 2007). Similarly, more than a dozen parallel losses of ribosomal protein (*rps*/*rpl*) gene have occurred in different lineages of land plants (Ruhlman and Jansen 2014). Three major pathways of gene loss have been detected in land plants: (1) gene transfer to the nucleus (*infA, rpl*22, *rpl*32 and *accD*), (2) substitution by a nuclear-encoded, mitochondrial targeted gene product (*rps*16), and (3) substitution by a nuclear-encoded protein for a plastid gene product (*accD, rpl*23) (Jansen and Ruhlman 2012). The multiple independent *ndh* gene loss in different lineages is supposed to belong to the third pathway (Ruhlman et al. 2015).

Selaginellaceae, one of the most ancient vascular plants with nearly 400 million years of evolutionary history (Banks 2009), is the largest family of lycophytes with ca. 750 species classified into the only genus *Selaginella* (Jermy 1990; Weststrand and Korall 2016b; Zhou et al. 2016). *Selaginella* species have highly diverse growth forms, including creeping, climbing, prostrate, erect and rosetted forms, and also inhabit an impressive range of habitats, from tropical rain forests to deserts, alpine and arctic habitats (Zhang et al. 2013). With such a high diversity in habitat and growth forms, complex plastomes with different structures are expected in Selaginellaceae (Tsuji et al. 2007). However, only four species of *Selaginella*, viz., *S. uncinata* (Tsuji et al. 2007), *S. moellendorffii* (Smith 2009), *S. tamariscina* (Xu et al. 2018), and *S. vardei* (Unpublished data), have been reported for their plastomes. Compared to the four species from Lycopodiaceae and Isoetaceae of lycophytes (Guo et al. 2016; Karol et al. 2010; Wolf et al. 2005; Zhang et al. 2017), plastomes of *Selaginella* are, indeed, far less conserved in both structures and gene contents. Both *S. uncinata* and *S. moellendorffii* belong to subg. *Stachygynandrum* based on both morphology-based classification (Jermy 1986) and a recent molecular-based classification (Weststrand and Korall 2016b). However, their plastomes show divergent variation in structure. Several rearrangements, such as a 20 kb fragment inversion, a 17 kb fragment transposition and gene duplications, exist in these two species (Smith 2009). The morphology of *S. tamariscina*, belonging to subg. *Stachygynandrum*, and *S. vardei*, belonging to subg. *Rupestrae* (sensu Weststrand and Korall 2016) (Weststrand and Korall 2016b) are also quite divergent from *S. uncinata* and *S. moellendorffii* in having rosetted plant and helically-arranged trophophylls, respectively. These two species both grow in extremely xeric habitat. Plastomes with DR rather than IR have been characterized in *S. tamariscina* (Xu et al. 2018) and *S. vardei* (Unpublished data) was characterized. Some other features of these plastomes are also quite distinctive. Only 9-12 different tRNA genes remain in *Selaginella*, and GC content in *Selaginella* plastomes is significantly higher (54.8% for *S. uncinata*, 51% for *S. moellendorffii*, 54.1% for *S. tamariscina*, and 53.2% for *S. vardei*) than the plastomes of other land plants (less than 43%) (Smith 2009; Tsuji et al. 2007). Such extensive rearrangement events and extraordinary gene content have never been reported in other lycophytes and fern families (Guo et al. 2016; Karol et al. 2010; Mower and Vickrey 2018).

The divergent variation in structure and gene content exhibited by Selaginellaceae plastomes make it an ideal family to study the complexity and diversity of plastomes. Furthermore, the extent of genomic change in other lineages of *Selaginella* species has not been fully investigated. Therefore, we sequenced a total of ten plastomes from species belonging to six different main lineages of Selaginellaceae using next generation sequencing and combined previously published plastomes of lycophytes to reach the following goals: 1) document plastome characteristics from major lineages of Selaginellaceae, 2) explore the evolutionary trajectory and dynamic conformations of DR/IR structure in plastomes of Selaginellaceae, and 3) reveal the potential correlations among plastome rearrangements, substitution rate, and number of repeats.

## Materials and Methods

### Taxon Sampling

Seven taxa (*S. lyallii, S. remotifolia, S. sanguinolenta, S. doederleinii, S. pennata, S. bisulcata*, and *S. hainanensi*s) from five main lineages of Selaginellaceae representing four subgenera of Zhou and Zhang (2015) and three subgenera of Weststrand and Korall (2016) were sampled (Table S1). The previously published plastomes of *S. tamariscina, S. moellendorffii*, and *S. uncinata* were re-sequenced to confirm their structures. Previously published plastomes of *S. vardei* (Unpublished data) and four outgroups from other lycophytes (*Huperzia lucidula* (NC_006861), *H. javanica* (KY609860), *H. serrata* (NC_033874) and *Isoetes flaccida* (NC_014675))(Guo et al. 2016; Karol et al. 2010; Wolf et al. 2005; Zhang et al. 2017) were used in the following analyses. We followed the classification of Zhou and Zhang (2015) to conveniently describe the lineages represented by our species.

### DNA Extraction, Sequencing and Assembly

The total genomic DNAs were isolated from silica-dried materials with a modified cetyl trimethylammonium bromide (CTAB) method (Li et al. 2013). Library construction was performed with NEBNext DNA Library Prep Kit (New England Biolabs, Ipswich, Massachusetts, USA) and sequencing was finished by Illumina HiSeq 2500 (Illumina, San Diego, California, USA). Illumina paired-end reads of each species were mapped to *S. uncinata* (AB197035) (Tsuji et al. 2007) and *S. moellendorffii* (FJ755183) (Smith 2009), with medium-low sensitivity in five to ten iterations in Geneious v. 9.1.4 (Kearse et al. 2012)(Biomatters, Inc., Auckland, New Zealand; https://www.geneious.com). The mapped reads were then assembled into contigs in Geneious. We also used bandage v. 0.8.1 (Wick et al. 2015), a program for visualizing de novo assembly graphs, to help select contigs of plastome and analyze de novo assemblies by importing the fastg file created by GetOrganelle (Jin et al. 2018). The contigs obtained from both ways were then combined and imported into Geneious to extend and assemble into the complete plastomes.

### Gene Annotation

Gene annotations were performed with local BLAST (Delsuc et al. 2005) using the plastomes of *H. serrata* (Guo et al. 2016), *I. flaccida* (Karol et al. 2010), *S. uncinata* (Tsuji et al. 2007), and *S. moellendorffii* (Smith 2009) as references. Putative start and stop codons were defined based on similarity with known sequences. The tRNAs were further verified using tRNAscan-SE version 1.21 (Lowe and Eddy, 1997; Schattner et al., 2005) and ARAGORN (Laslett and Canback 2004). Circular and linear plastome maps were drawn in OGDraw version 1.2 (Lohse et al., 2007).

### PCR Confirmation of the Plastome Structure of Representative Species in Four Subgenera

We selected eighteen representatives (Table S2) from four subgenera to confirm whether the DR structure is ubiquitous in plastomes of three subgenera and IR structure only exists in the derived subgenus. Primers were designed at the flank of rearrangement end points and DR/IR region boundaries. Considering the distant relationship among these subgenera, we designed the primers (Table S3) for each subgenus, respectively. The PCR amplifications were performed in a total volume of 25μL containing 2.5 μL of 10X Ex Taq ™ Buffer, 2.5 μL of dNTP Mixture (2.5 mM each), 2 μL of each primer (5 mM), 0.15 μL of TaKaRa Ex Taq ™ (5 units/µl) and 20 ng of template DNA. Cycling conditions were 94 °C for 3 min, followed by 35 cycles of 94 °C for 30 s, 52 °C for 1 min and 72 °C for 1.5 min, and a final extension of 72 °C for 10 min. The PCR products were verified by electrophoresis in 1% agarose gels stained with ethidium bromide and sequenced by the Company of Majorbio, Beijing, China.

### Plastome Rearrangement Analyses

In order to identify the putative presence of large structural variation within the *Selaginella* plastomes, whole genome alignment among the 14 lycophyte species (twelve Selaginellaceae species, *I. flaccida*, and *H. serrata*) was performed using the progressiveMauve algorithm in Mauve v 2.3.1 (Darling et al. 2010). A copy of DR/IR was removed from the plastomes. The Locally colinear blocks (LCBs) identified by the Mauve alignment were numbered to estimate the genome rearrangement distances (Table S4). Genes in each block were also listed (Table S5). Two types of genome rearrangement distances, break point (BP) and inversions (IVs), were calculated using the web serve of the Common Interval Rearrangement Explore (CREx) (Bernt et al. 2007) using the conserved plastome of *H. serrata* as a reference.

### Repeats Analyses

Repeats within the 14 lycophytes plastomes were identified by RepeatsFinder (Volfovsky et al. 2001) with default parameters (repeat size >15 bp). One copy of the DR/IR was removed from all plastomes used. The circular layouts of repeats in plastome were visualized using the *circlize* package (Gu et al. 2014) in R. Furthermore, the correlation between the number of repeats and the degree of genome rearrangements, BPs and IVs distance, were tested using Pearson test in R v. 3.4.1 (R Development Core Team2012).

### Nucleotide Substitution Rate Analyses

Forty-six protein-coding genes (Table S6) from single copy regions of 12 Selaginellaceae species and one outgroup, *I. flaccida*, were extracted from the plastomes and aligned at the protein level by MAFFT (Katoh and Standley 2013) using the translation-aligned function in Geneious v. 9.1.4 (Kearse et al. 2012). Poorly aligned regions were removed by using Gblocks v. 0.91b (Castresana 2000) with default parameters. The dataset for substitution rate comparison between plastomes with DR and IR/DR-coexisting structure in Selaginellaceae includes all twelve Selaginella species. The dataset for comparison of genes inside or outside the ca. 50 kb inversion (Table S6) includes eight *Selaginella* species with plastomes of DR structure. The pairwise synonymous substitution rate (d*S*), nonsynonymous substitution rate (d*N*), and d*N*/d*S* of each individual gene was estimated using PAML v. 4.9 (run mode=-2) (Yang 2007) with codon frequencies determined by the F3 × 4 model. The significance of differences of d*S*, d*N*, and d*N*/d*S* was assessed using Wilcoxon rank sum tests in R v. 3.4.1 (R Development Core Team2012).

### Phylogenetic Analysis and Divergence Time Estimation

Forty-six protein-coding genes (Table S6) with 21,382 bases shared by 16 lycophyte species (twelve *Selaginella*, one *Isoetes* and three *Huperzia*) were extracted and aligned using Multiple Alignment in Geneious v. 9.1.4 (Kearse et al. 2012) under the automatic model selection option with some manual adjustments. The 1^st^ and 2^nd^ sites of each codon were selected in MEGA 7.0 (Tamura et al. 2007) to eliminate the effect of 3^rd^ site base substitution saturation. Phylogenetic analysis was performed using maximum likelihood methods on the RAxML web server with 1000 bootstrap replicates and the GTR+G model was selected based on Akaike information criterion (AIC) in jModeltest 2.1.7 (Darriba et al. 2012). The divergence time of DR/IR occurrence was estimated using BEAST version 1.8.2 (Suchard et al. 2018) with two fossil calibration nodes employed. A fossil calibration of the root age corresponded to the split of Lycopodiopsida and Isoetopsida (Figure 4, node A: [392-451 Ma]) (Morris et al. 2018) with a selection of normal prior distributions. The other fossil calibration of the node separated Isoetaceae and Selaginellaceae (Figure 4, node B: [3722-392 Ma]) (Kenrick and Crane 1997) with a lognormal prior distribution. A relaxed clock with lognormal distribution of uncorrelated rate variation was specified, and a birth-death speciation process with a random starting tree was adopted. The MCMC chain was run for 500 million generations, sampling every 1000 generations. The effective sample size (ESS) was checked in Tracer v 1.5 (Rambaut and Drummond 2009). The maximum clade credibility tree was generated using TreeAnnotator in BEAST and the tree was plotted using FigTree v. 1.4.3 (Rambaut 2017). The events of DR/IR originate, rearrangements, and DR/IR expansion/contraction in Selaginellaceae was mapped on the phylogenetic tree to explore the evolutionary trajectory of Selaginellaceae plastomes.

## Results

### Characteristics of Selaginellaceae Plastomes

The general features of twelve Selaginellaceae plastomes and four other lycophytes are summarized in Table 1 and Table S1. Compared to other lycophytes, the plastomes of Selaginellaceae showed remarkable variation in size, ranging from roughly 110 kb in *S. lyallii* to 147 kb in *S. sanguinolenta*. Size variability was partly due to the IR, which expanded to ca. 16 kb in *S. sanguinolenta* and reduced to ca. 10 kb in *S. lyallii*. Gene content was also variable in Selaginellaceae plastomes due to a number of gene losses (Table 1, Figure 1). Lycophyte plastomes generally contained 120 different genes (86 protein-coding genes, 30 tRNA genes and 4 *rrn* genes), whereas it ranged from 73 different genes (61 protein-coding genes, 8 tRNA genes and 4 *rrn* genes) in *S. tamariscina* to 102 different genes (76 protein-coding genes, 22 tRNA genes and 4 *rrn* genes) in *S. sanguinolenta*. Intron loss was detected in *atpF, clpP, rpo*C1, and *ycf*3 genes. The GC content in Selaginellaceae plastomes was significantly higher than those in Isoetaceae and Lycopodiaceae. The average GC content was 53.2% in Selaginellaceae (ranging from 50.7% in *S. lyallii* to 56.5% in *S. remotifolia*) and 36.7% in other lycophytes (Table 1).

**Table 1:** Plastome Characteristics for Representative Selaginellaceae in Comparison to Four Lycophytes

| Species | Size<br>(bp) | LSC (bp) | SSC (bp) | IR (bp) | No.<br>different<br>genes | No. different protein-coding genes<br>(duplicated) | No. different<br>tRNA genes<br>(duplicated) | No. different<br>rRNA genes<br>(duplicated) | GC<br>content<br>(%) |
| --- | --- | --- | --- | --- | --- | --- | --- | --- | --- |
| *S. uncinata* | 144,161 | 77,752 | 40,851 | 12,779 | 97 | 81 (3) | 12 (3) | 4 (4) | 54.9 |
| *S. hainanensis* | 144,201 | 77,780 | 40,819 | 12,801 | 97 | 81 (3) | 12 (3) | 4 (4) | 54.8 |
| *S. bisulcata* | 140,509 | 55,598 | 59,659 | 12,626 | 94 | 78 (3) | 12 (3) | 4 (4) | 52.8 |
| *S. pennata* | 138,024 | 54,979 | 59,847 | 11,599 | 90 | 77 (3) | 9 (2) | 4 (4) | 52.9 |
| *S. moellendorffii* | 143,525 | 58,198 | 61,129 | 12,099 | 96 | 78 (1) | 14 (2) | 4 (4) | 51 |
| *S. doederleinii* | 142,752 | 57,841 | 62,865 | 11,023 | 96 | 78 (1) | 14 (2) | 4 (4) | 51.1 |
| *S. tamariscina* | 126,700 | 53,299 | 47,741 | 12,830 | 73 | 61 (1) | 8 (1) | 4 (4) | 54.1 |
| *S. sanguinolenta* | 147,148 | 54,436 | 59,650 | 16,531 | 102 | 76 (4) | 22 (3) | 4 (4) | 50.8 |
| *S. remotifolia* | 131,866 | 46,351 | 55,851 | 14,832 | 87 | 71 (3) | 12 (2) | 4 (4) | 56.5 |
| *S. lyallii* | 110,411 | 44,943 | 45,276 | 10,096 | 76 | 60 (1) | 13 (2) | 4 (4) | 50.7 |
| *S. indica* | 122,460 | 45,711 | 48,395 | 14,177 | 78 | 62 (3) | 12(3) | 4 (4) | 53.6 |
| *S. vardei* | 121,254 | 45,792 | 47,676 | 13,893 | 78 | 62 (3) | 12(3) | 4 (4) | 53.2 |
| *I. flaccida* | 145,303 | 91,862 | 27,275 | 13,118 | 123 | 87 (3) | 32 (5) | 4 (4) | 37.9 |
| *H. javanic* | 154,415 | 104,120 | 19,667 | 15,314 | 119 | 86 (3) | 29 (4) | 4 (4) | 36.4 |
| *H. lucidula* | 154,373 | 104,088 | 19,657 | 15,314 | 119 | 86 (3) | 29 (4) | 4 (4) | 36.2 |
| *H. serrata* | 154,176 | 104,080 | 19,658 | 12,219 | 120 | 86 (3) | 30 (5) | 4 (4) | 36.3 |

**Figure 1.**
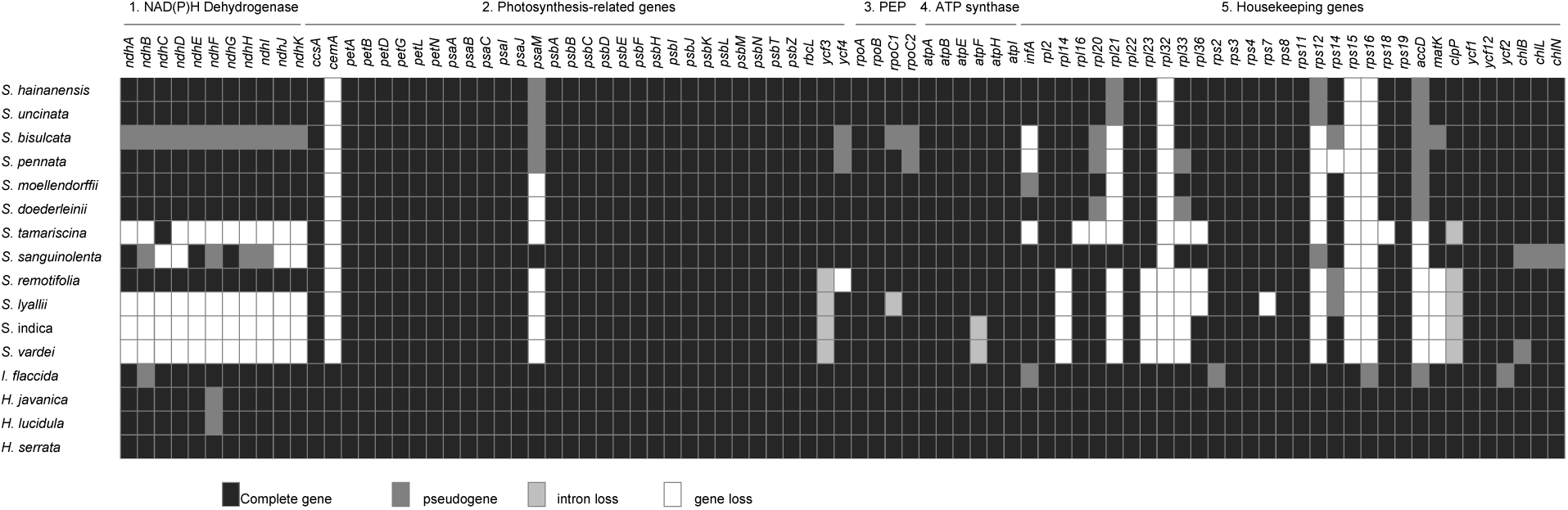
Protein-coding genes in Selaginellaceae and other lycophytes. Intact genes per species are indicated by black boxes, dark grey box represent putative pseudogenes, light grey and white boxes mark intron and gene losses, respectively. PEP—plastid-encoded RNA polymerase.

A noticeable plastome structure with DR was documented in *S. tamariscina* (Xu et al. 2018) and *S. vardei* (Unpublished data). With our newly selected species being sequenced, we assembled the plastomes of seven species (*S. lyallii, S. remotifolia, S. sanguinolenta, S. doederleinii, S. moellendorffii, S. bisulcata*, and *S. pennata*) into the DR structure (Figure 2). Followed with the DR structure, we found that the length of two single copy regions (LSC and SSC) changed into almost equal size. However, the length of LSC (44.9 kb – 57.8 kb) was slightly shorter than that of SSC (45.3 – 62.8 kb) (Table 1), which mainly resulted from a relocation of ca. 35 kb fragments from LSC to SSC region. Only the plastomes of *S. hainanensis* and *S. uncinata* were assembled into the typical IR structure (Figure 2). We consider the assembled plastome structures as master forms in the following analyses.

**Figure 2.**
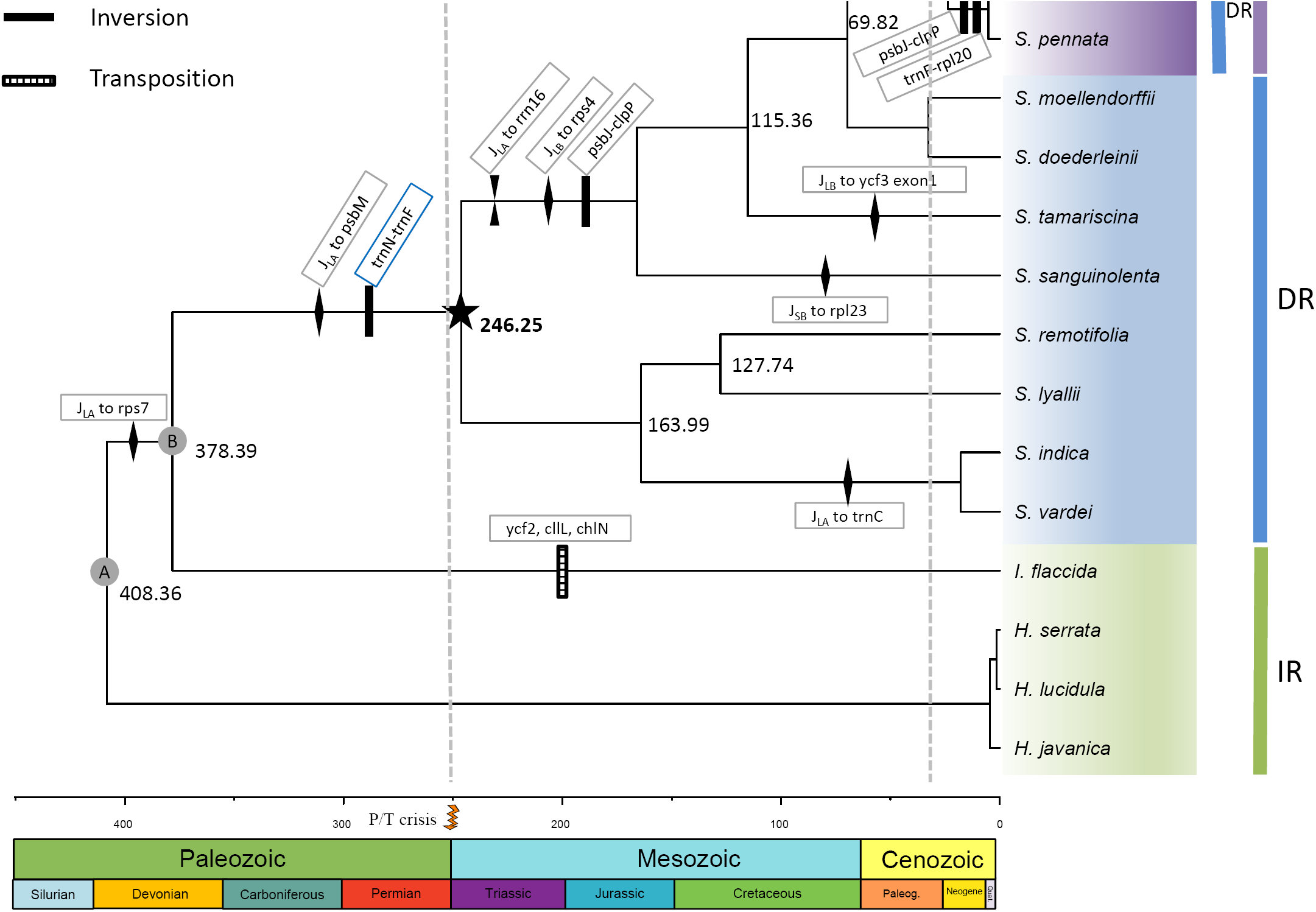
Phylogeny reconstruction and time divergence estimation of lycophytes with plastome rearrangement events mapped on the branches. Node A-B represent the calibration nodes. Node A: fossil calibration of the root age corresponding to the split of Lycopodiopsida and Isoetopsida (392-451 Ma); node B: fossil node separating Selaginellaceae and its sister family Isoetaceae (372-392 Ma). Black star represents the occurrence time of plastomes with DR structure, and grey star represents the occurrence time of plastomes with IR/DR-coexisting structure.

### Confirmation of DR/IR Structures in Plastomes of Representative Species

The DR structure in plastome of *S. vardei* for subg. *Rupestrae sensu* Weststrand and Korall (2016) (*Selaginella* sect. *Homoeophyllae sensu* Zhou and Zhang (2015)), have been confirmed by Zhang et al. (2018). The PCR confirmation of eighteen representative species with different structure (Table S2) from four subgenera *sensu* Zhou and Zhang (2015) suggested that DR structure are ubiquitous in subg. *Stachygynandrum*, subg. *Pulviniella*, subg. *Ericetorum*, and subg. *Boreoselaginella* (Figure S2 **B**, **C**, **D**). Particularly, the re-sequenced plastome of *S. moellendorffii* was also found to possess DR structure, which is quite inconsistent with the previously published IR structure (Smith 2009). The PCR confirmation of nine species from the same subgenus as *S. moellendorffii* further supported the DR structure (Figure S2 **B**). Therefore, we followed the re-sequenced DR-possessing plastome of *S. moellendorffii* for the following analyses. For subg. *Heterostachys*, the PCR confirmation of five species indicated that the IR structure only existed in the sect. *Oligomacrosporangiatae* which *S. uncinata* and *S. hainanensis* were located (Figure S2 **A**).

### Gene/Intron Loss and Pseudogenes

Gene loss or pseudogenization is a remarkable character of plastomes of Selaginellaceae including tRNA gene loss, protein-coding gene loss and intron loss.

### tRNA Gene Loss

The most noticeable feature is the tRNA gene loss (Table 1, Figure 3). Plastomes in other land plants usually contain 30 different tRNA genes while Selaginellaceae plastomes have experienced an extensive tRNA gene loss and varies greatly in different lineages. Twenty-two different tRNA genes were annotated in plastome of *S. sanguinolenta*, whereas only eight different tRNA genes existed in *S. tamariscina* plastome. Some vestiges of tRNA genes (e.g. *trnA-UGC* exon2 and *trnI-GAU* exon1 between *rrn*16 and *rrn*23, *trnV-UAC* exon1 between *trnM-CAU* and *trnF-GAA*) were observed in plastome of *S. sanguinolenta*.

**Figure 3.**
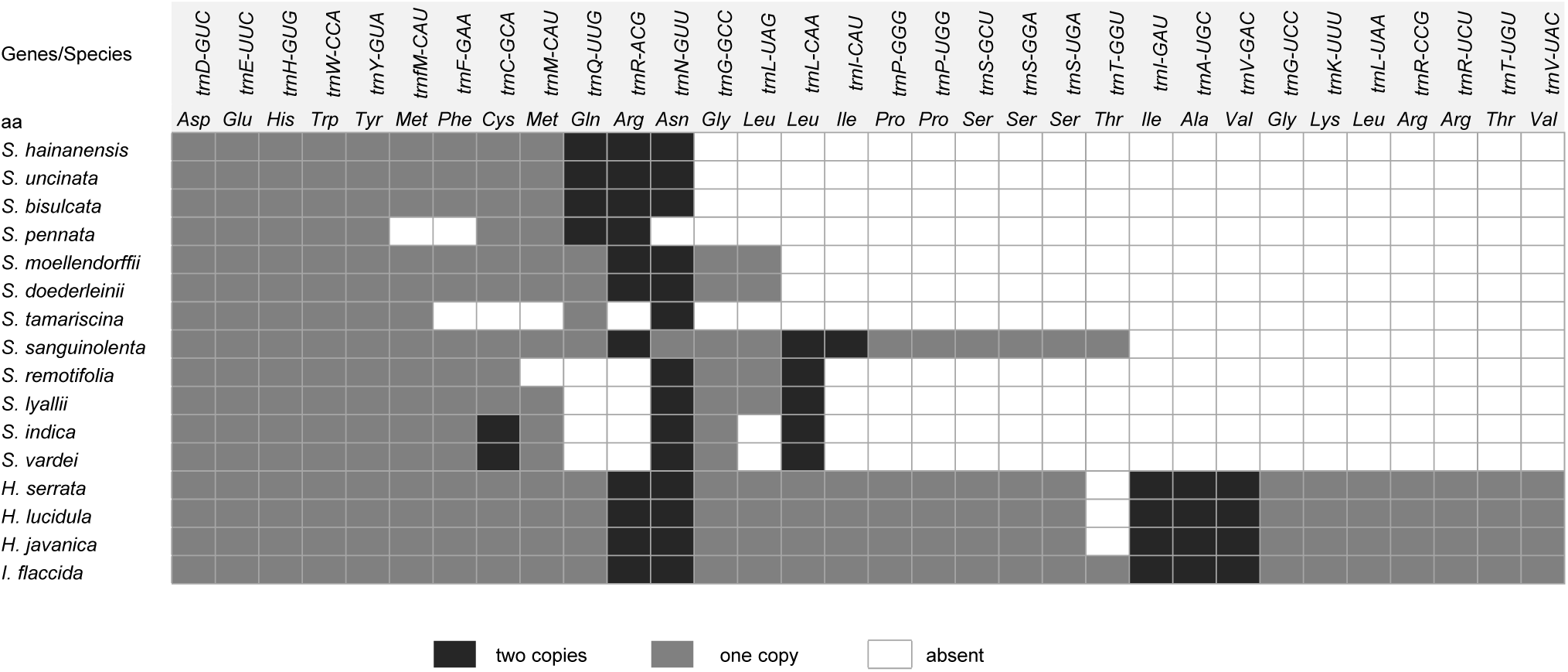
tRNA genes in Selaginellaceae and other lycophytes. Black box represents tRNA genes with two copies, grey box represents tRNA genes with one copy, and white box represents tRNA gene loss.

**Figure 4.**
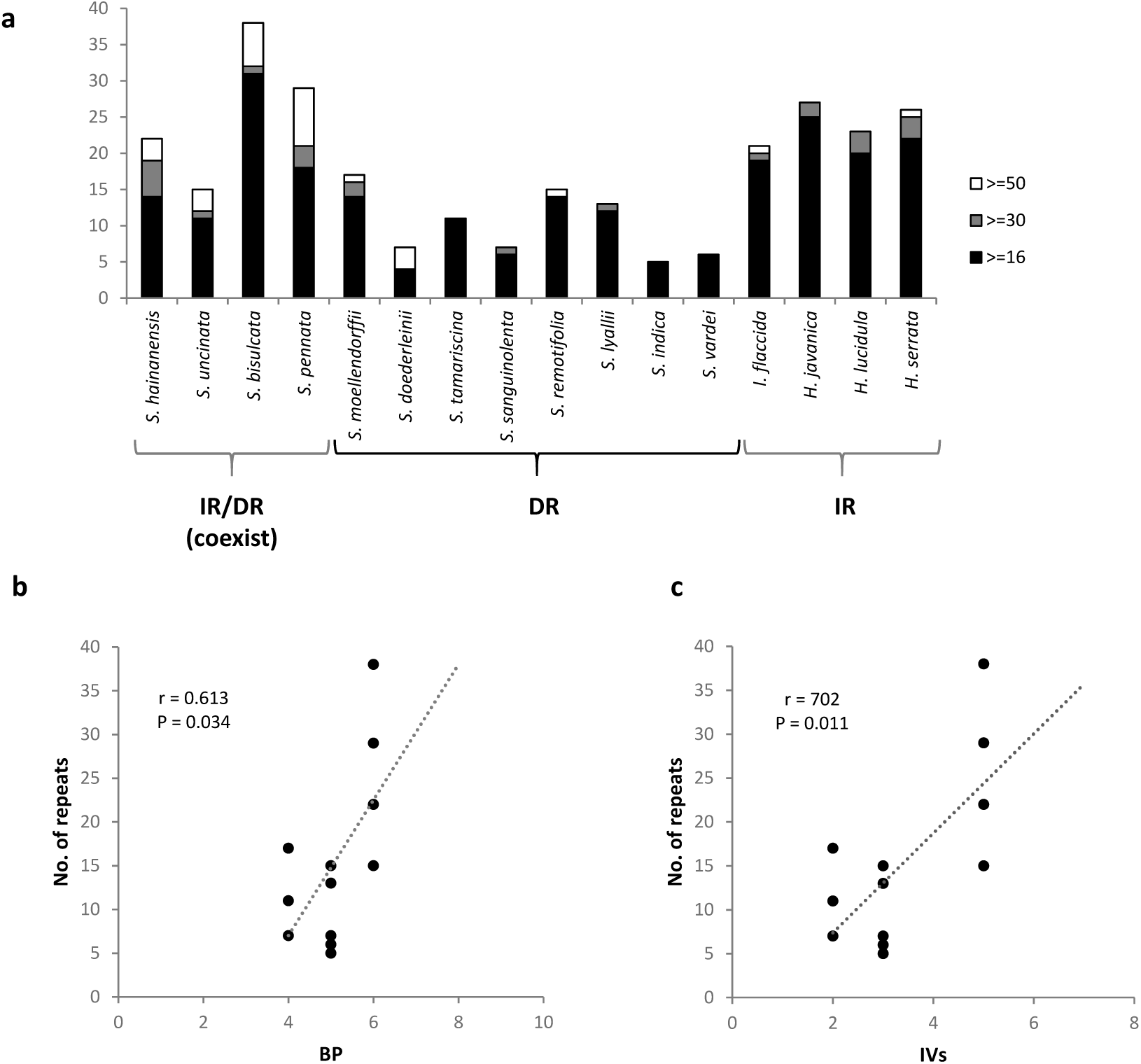
Statistics of repeat analyses in Selaginellaceae. **a**, The number of different size repeats; **b,** Correlation of BP distance with the number of repeats; **c,** Correlation of IV distance with the number of repeats.

### Protein-coding Gene/Intron Loss

The loss and putative pseudogenization (with internal stop codons) of protein-coding genes in Selaginellaceae plastomes are shown in figure 1, mainly focusing on NAD(P)H-dehydrogenase complex-encoding genes (*ndh* genes) and ribosomal proteins-encoding genes (*rpl*/*rps* genes). In *Selaginella*, the gene loss or pseudogenization of *ndh* genes occurred to a different extent. All *ndh* genes were lost in *S. vardei, S. indica*, and *S. lyallii*, and only one functional *ndhC* remained in *S. tamariscina*, whereas in *S. sanguinolenta*, four *ndh* genes (B, F, H, I) became pseudogenes, four (C, D, J, K) were completely lost, and other three (A, E, G) were still functional. The whole set of n*dh* genes in *S. bisulcata* were all pseudogenized because of internal stop codons caused by reading frame shift, whereas they were intact and functional in its sister species, *S. pennata*. In addition to the whole set of gene loss in specific species, several genes were lost across the whole family. The genes of *cemA, rpl*32, *rps*15, and *rps*16 were absent in plastomes of all *Selaginella* species, whereas present in all four outgroups from *Isoetes* and *Huperzia*. The gene *accD* was non-functional (pseudogenized or lost) in all Selaginellaceae and *I. flaccida*, but was functional in Lycopodiaceae. Besides, a number of ribosomal genes (*rpl*14, 16, 20, 21, 23, 33, 36 and *rps*12, 14, 18) were lost or pseudogenized independently in different lineages of Selaginellaceae. Although *infA* is present in all bryophyte and fern lineages, it is pseudogenized in the plastomes of *I. flaccida* and *S. moellendorffii* and lost in three *Selaginella* species (*S. bisulcata, S. pennata*, and *S. tamariscina*). *MatK* gene was absent in *S. remotifolia, S. lyallii, S. indica* and *S. vardei*, and pseudogenized in *S. bisulcata*. *RpoC*1 and *rpoC*2 were pseudogenized in *S. bisulcata* and *rpoC*1 was pseudogenized in *S. pennata*. All three *chl* genes were pseudogenized in *S. sanguinolenta*, and only *chlB* was pseudogenized in *S. vardei*. However, they were all intact in other *Selaginella* species. The gene *psaM* was non-functional (pseudogenized or lost) in all *Selaginella* species except *S. sanguinolenta*, which possessed an intact *psaM* gene. Another event worth noticing was the intron loss in Selaginellaceae. Two introns remained in *clpP* of *S. doederleinii* and *S. sanguinolenta*, none in *S. tamariscina, S. remotifolia, S. lyallii, S. indica* and *S. vardei*, and one remained in other species of *Selaginella*. The intron of *atpF* gene was lost in *S. indica* and *S. vardei*, the intron of *rpoC1* was lost in *S. lyallii*, and the intron 2 of *ycf*3 gene was lost in *S. vardei, S. indica, S. lyallii* and *S. remotifolia*.

### Rearrangement Events in Plastomes of Selaginellaceae

Nineteen locally collinear blocks (LCBs) (Table S5) shared by Selaginellaceae and outgroups were identified by Mauve whole plastome alignment. Each LCB for 12 plastomes was numbered from 1 to 19 and assigned a ± showing strand orientation (Table S4). The order of the 19 LCBs in each plastome was number coded for estimating breakpoint (BP) and inversions (IVs) distances between plastomes. The pairwise comparison of the two types of plastome rearrangement distances is shown in Table 2. The two distances were highly correlated (P <0.001, r = 0.98). Both distances were used as the estimation of the degree of genome rearrangement in the later analyses. The plastome rearrangement between *I. flaccida* and *S. vardei* was described in Zhang et al. (2018). The plastomes of *S. vardei, S. indica, S. lyallii, S. remotifolia* and *S. sanguinolenta* were almost syntenic except the different extent loss of *ndh* genes (Figure 1, Figure S1) and slightly change of DR boundaries (Figure 2). Compared with the plastome of *S. sanguinolenta*, an inversion of 8 kb *psbJ*-*clpP* fragment (block -15, - 14) was observed in the plastome of *S. tamariscina*. The *psbJ*-*clpP* inversion was also shared by the plastomes of *S. doederleinii* and *S. moellendorffii*, which are basically snytenic with that of *S. tamariscina*. The inversion of a ca. 20 kb *trnC*-*psbI* fragment (block 4, 5), absent in *S. moellendorffii* and other early diverged species, showed a derived character in the plastomes of *S. bisulcata, S. pennata, S. hainanensis*, and *S. uncinata*. Two extra inversions existed when comparing the plastomes of *S. moellendorffii* and *S. bisulcata*. The first inversion of a 9 kb *psbJ*-*clpP* fragment (block 14, 15) was located at one end of the second inversion of 20 kb *rpl*20-*trnF* fragment (block -14, -13, -12) with *clpP* (block -15) being excluded. The plastome organization of *S. pennata* was basically syntenic with its sister species, *S. bisulcata*.

**Table 2.**
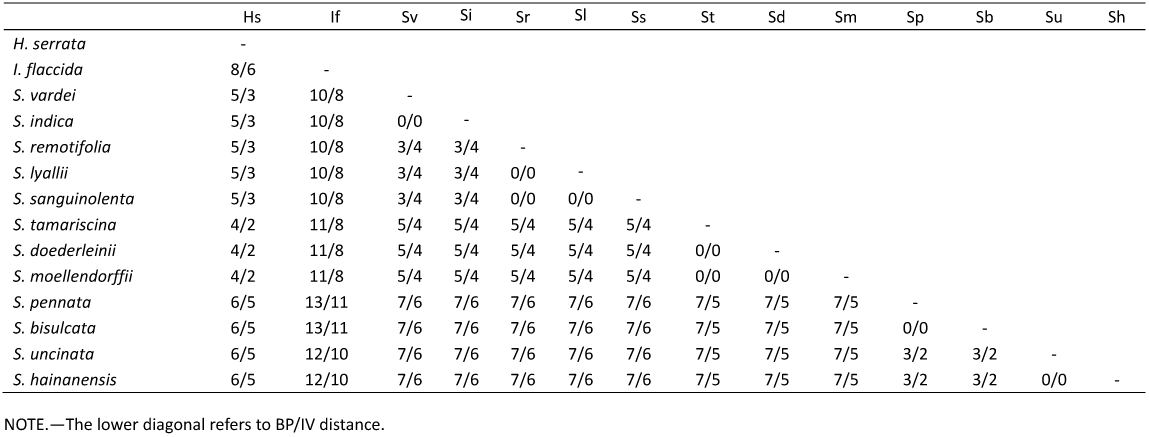
Pairwise Comparison of Genome Rearrangement Distances.

|  | Hs | If | Sv | Si | Sr | Sl | Ss | St | Sd | Sm | Sp | Sb | Su | Sh |
| --- | --- | --- | --- | --- | --- | --- | --- | --- | --- | --- | --- | --- | --- | --- |
| *H. serrata* | - |  |  |  |  |  |  |  |  |  |  |  |  |  |
| *I. flaccida* | 8/6 | - |  |  |  |  |  |  |  |  |  |  |  |  |
| *S. vardei* | 5/3 | 10/8 | - |  |  |  |  |  |  |  |  |  |  |  |
| *S. indica* | 5/3 | 10/8 | 0/0 | - |  |  |  |  |  |  |  |  |  |  |
| *S. remotifolia* | 5/3 | 10/8 | 3/4 | 3/4 | - |  |  |  |  |  |  |  |  |  |
| *S. lyallii* | 5/3 | 10/8 | 3/4 | 3/4 | 0/0 | - |  |  |  |  |  |  |  |  |
| *S. sanguinolenta* | 5/3 | 10/8 | 3/4 | 3/4 | 0/0 | 0/0 | - |  |  |  |  |  |  |  |
| *S. tamariscina* | 4/2 | 11/8 | 5/4 | 5/4 | 5/4 | 5/4 | 5/4 | - |  |  |  |  |  |  |
| *S. doederleinii* | 4/2 | 11/8 | 5/4 | 5/4 | 5/4 | 5/4 | 5/4 | 0/0 | - |  |  |  |  |  |
| *S. moellendorffii* | 4/2 | 11/8 | 5/4 | 5/4 | 5/4 | 5/4 | 5/4 | 0/0 | 0/0 | - |  |  |  |  |
| *S. pennata* | 6/5 | 13/11 | 7/6 | 7/6 | 7/6 | 7/6 | 7/6 | 7/5 | 7/5 | 7/5 | - |  |  |  |
| *S. bisulcata* | 6/5 | 13/11 | 7/6 | 7/6 | 7/6 | 7/6 | 7/6 | 7/5 | 7/5 | 7/5 | 0/0 | - |  |  |
| *S. uncinata* | 6/5 | 12/10 | 7/6 | 7/6 | 7/6 | 7/6 | 7/6 | 7/5 | 7/5 | 7/5 | 3/2 | 3/2 | - |  |
| *S. hainanensis* | 6/5 | 12/10 | 7/6 | 7/6 | 7/6 | 7/6 | 7/6 | 7/5 | 7/5 | 7/5 | 3/2 | 3/2 | 0/0 | - |

NOTE.—The lower diagonal refers to BP/IV distance.

The plastomes of *S. bisulcata* and *S. pennata* were assembled into DR structure whereas the plastomes of both *S. uncinata* and *S. hainanensis*, which belong to the same subgenus, retained the typical IR structure as observed in other land plants. Therefore, how the DR region change back to IR again is an intriguing phenomenon. The comparison between plastomes of *S. bisulcata* and *S. uncinata* displayed two inversions. The first inversion was a 20 kb fragment of *rpl*20-*trnF* (block 12, 13, 14); followed by the inversion of 65 kb fragment of *trnD*-*rpl*20 (block 9, 10, 11, -17, -16, - 15, -14, -13, -12). The first inversion occurred at one end of the second inversion. With one copy of repeat region (DR_B_) inside, the second inversion changed *S. uncinata* repeat regions into an IR structure. Since *S. hainanensis* is sister to *S. uncinata*, the plastome structure is basically syntenic with each other.

### Expansion and Contraction of DR/IR Regions in *Selaginella* Plastomes

Repeat regions of plastomes in different lineages of *Selaginella* ranged from 10,096 bp in *S. lyallii* to 16,531 bp in *S. sanguinolenta* (Table 1). In the early diverged lineages of *S. vardei, S. indica, S. lyallii, S. remotifolia* and *S. sanguinolenta*, one end of the DR_B_ region expanded and incorporated genes (*rps*7, *ndhB, psbM, petN* and *trnC*) from LSC region to a different extant, whereas the other end of the DR_B_ region contracted to *trnN* or *trnR* (Figure 2, Figure S1). In the derived lineages like *S. tamariscina, S. doederleinii, S. bisulcata* and *S. hainanensis*, one end of the DR_A_ next to the LSC region contracted to *rrn*16 or *rpl*23 whereas the other end of the DR_A_ next to the SSC region expanded to include *rps*4 and even one exon of *ycf*3 (Figure 2, Figure S1).

### Repeats in *Selaginella* Plastomes

The total number of repeats in Selaginellaceae plastomes were slightly lower than that in Isoetaceae and Lycopodiaceae plastomes, whereas repeats with the size above 50 bp were mostly in Selaginellaceae plastomes (Figure 4A, Figure S2). Among the sixteen plastomes compared, *S. bisulcata* contained the most repeats (38) in lycophytes, with *S. bisulcata* and *S. pennata* possessing the most repeats above 50 bp (8), whereas *S. indica* had the fewest (5) in total. The number of repeats varied among the species of Selaginellaceae, and the plastomes of *S. bisulcata* and *S. pennata* contained the most repeats (38 and 29) of all. The degree of plastome rearrangement estimated by BP and IV distances were moderately correlated with the total number of repeats (BP, P=0.034, r =0.613; IVs, P=0.011, r=0.702) (Figure 4B, C) in all twelve *Selaginella* species, but were not correlated in the plastomes with DR structure (BP, P=0.498; IVs, P=0.498, data not shown). We did find some repeats, which are able to mediate homologous recombination, flanking several rearrangement end points. A pair of short repeats existed at the flank of ca. 20 kb *psbI*-*trnC* inversion in *S. bisulcata* (48 bp) and *S. pennata* (60 bp) (Figure S2C, D), and another pair of repeats at the flank of 10 kb *psbJ*-*clpP* inversion in *S. doederleinii* (166 bp) and *S. moellendorffii* (264 bp) (Figure S2E, F) suggesting that repeats larger than 50 bp may have facilitated rearrangements in Selaginellaceae. Besides, a pair of ca. 1.8 kb inverted repeats exists in plastomes of *S. bisulcata* and *S. pennata*, and a pair of ca. 2.7 kb direct repeats exists in plastomes of *S. uncinata* and *S. hainanensis*. These two pairs of repeats, together with DR and IR, are able to frequently mediate diverse homologous recombination, and create approximately equal stoichiometric subgenomes and isomers. Therefore, both IR and DR structure exist dynamically (IR/DR coexist) in plastomes of *S. bisulcata, S. pennata, S. uncinata*, and *S. hainanensis*. (Figure 6).

**Figure 5.**
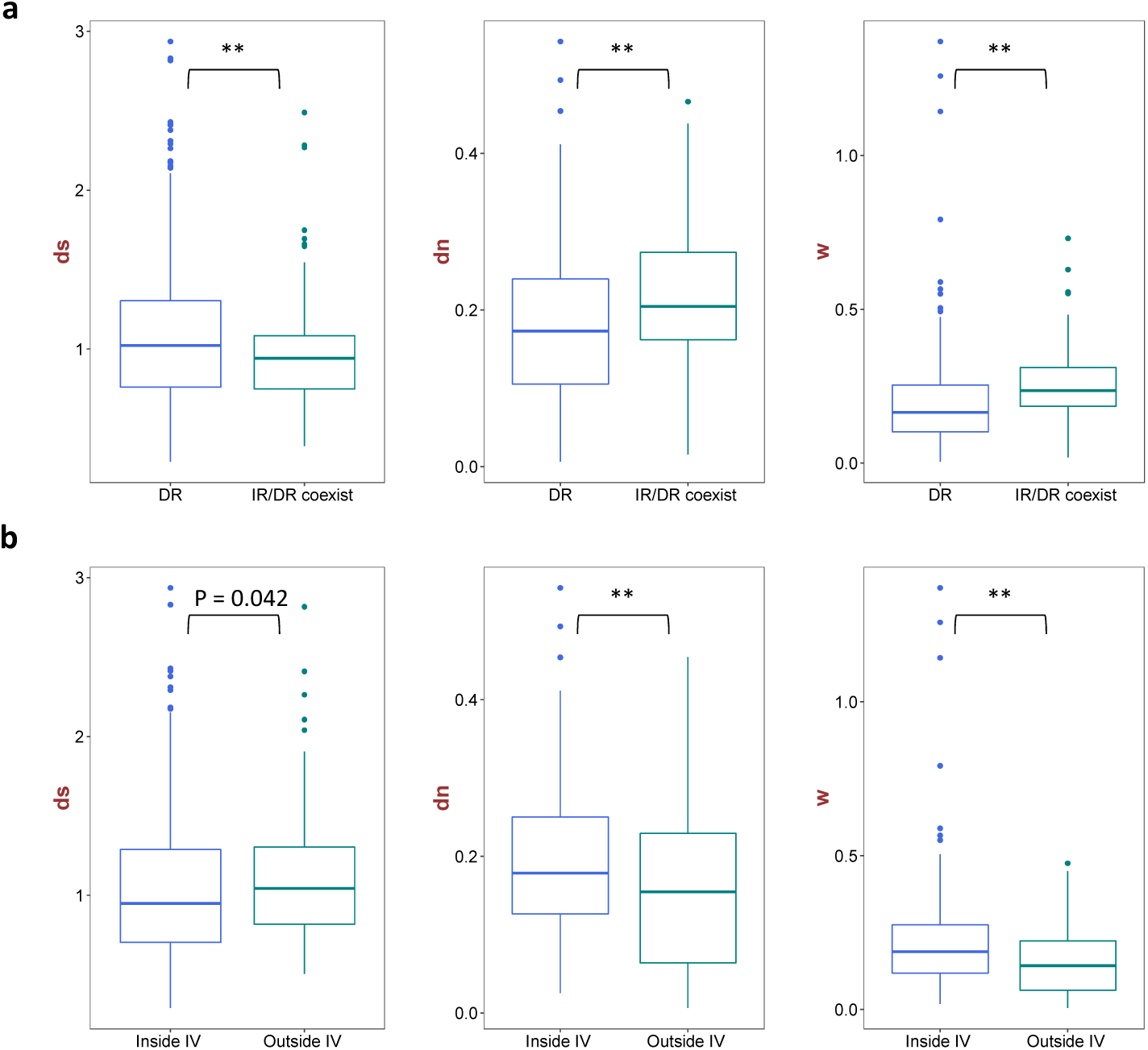
Nucleotide substitution rate analyses in Selaginellaceae. **a**, Substitution rate of genes from DR-possessing plastomes and IR/DR-coexisting plastomes; **b,** Substitution rate of genes inside IV and outside IV from DR-possessing plastomes.

**Figure 6.**
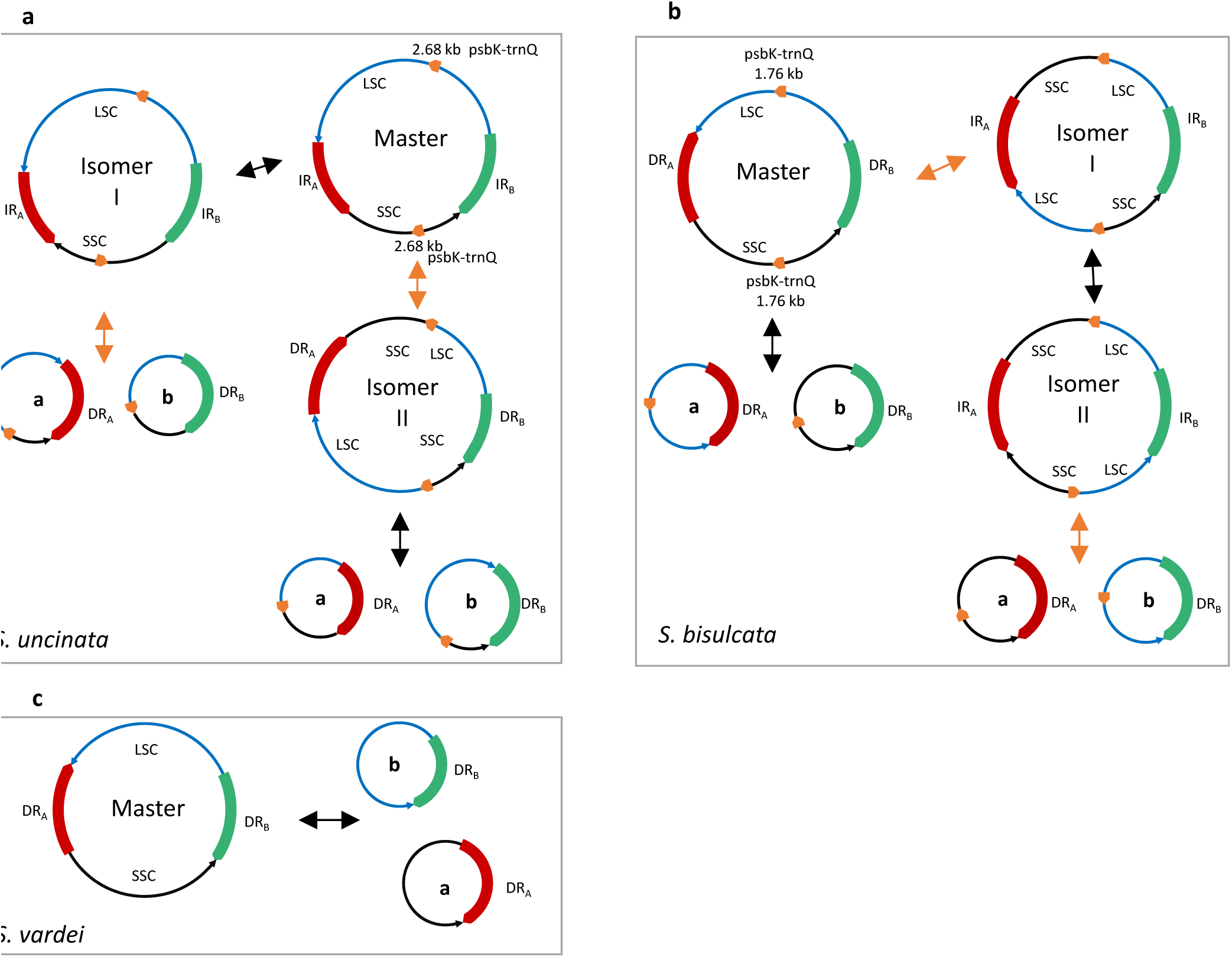
Dynamic structures of plastomes with DR/IR structure of Selaginellaceae. **a**, Recombination activities between repeats in plastomes of *S. uncinata* and *S. hainanensis*. The red and green block represents two copies of IR with arrows at the end showing orientation and the short orange blocks represent two copies of repeats (*psb*K-*trn*Q). The black double arrows show recombination between IR and the orange double arrows show recombination between repeats *psb*K-*trn*Q; **b,** Recombination activities between repeats in plastomes of *S. bisulcata* and *S. pennata*. The red and green blocks represent two copies of DR with arrows at the end showing orientation and the short orange blocks represent two copies of repeats (*psb*K-*trn*Q). The black double arrows show recombination between DR and the orange double arrows show recombination between repeats *psb*K-*trn*Q; **c,** Recombination activities between repeats in plastomes with DR structure.

### Nucleotide Substitution in *Selaginella* Plastomes

The pairwise substitution rate comparison of 46 genes from single copy regions of twelve *Selaginella* plastomes showed that the d*S* values for the genes of DR-possessing plastomes were significantly higher than those of IR/DR-coexisting plastomes (P<0.01), whereas the opposite trend was observed for the d*N* value and d*N*/d*S* (P<0.01, P<0.01) (Figure 5). Of the 46 genes, 28 genes were inside the ca. 50 kb inversion and 18 genes were outside the inversion for DR-possessing plastomes. Comparison between genes inside and outside IV showed that d*S* was slightly lower in genes inside IV than outside IV without significant difference (P=0.042), whereas the opposite trend was observed for d*N* and d*N*/d*S* (P<0.01, P<0.01) (Figure 5).

### Phylogenetic Reconstruction and Divergence Time Estimation

Phylogenetic relationships of 46 protein-coding genes within Selaginellaceae were basically congruent with the previously published results (Weststrand and Korall 2016a; Zhou et al. 2016) except the phylogenetic position of *S. sanguinolenta* (Figure 2). *Selaginella sanguinolenta* was placed in the second earliest diverging subg. *Boreoselaginella* and was sister to the rest of the genus except subg. *Selaginella* in Zhou et al. (2016). In Weststrand and Korall (2016a), *S. sanguinolenta* was found in two different positions (position α and position β), and the phylogenetic position of *S. sanguinolenta* in our newly reconstructed phylogeny was congruent with the position β with 100% support. The split between Selaginellaceae and its sister family Isoetaceae occurred at 375 million years ago (Ma) (Kenrick and Crane 1997). We inferred that the occurrence time of the DR structure and the recurrence time of the IR structure (IR/DR-coexisting) was at 246 Ma and 23 Ma, respectively (Figure 2). The DR structure showed the early diverged state whereas IR structure was the derived one in the most evolved lineage in the family Selaginellaceae when mapped on the phylogenetic tree (Figure 2). The 20 kb *trnC*-*psbI* inversion showed a derived character for *S. pennata, S. bisulcata, S. hainanensis* and *S. uncinata*. The shared 9 kb *psbJ*-*clpP* inversion among *S. tamariscina, S. doederleinii*, and *S. moellendorffii* suggested their relatively close relationship (Figure 2).

## Discussion

### Gene/Intron Losses and Pseudogenes

Compared with *S. uncinata* (Tsuji et al. 2007) and *S. moellendorffii* (Smith 2009), the tRNA loss was even severe in the basal *Selaginella* species with only eight different tRNA genes remaining in *S. tamariscina*. However, 22 tRNA genes were annotated in the plastome of *S. sanguinolenta* (Table 1, Figure 3) with the presence of three “vestigial” tRNA genes, which were also found in *S. uncinata* plastomes (seven “vestigial” tRNA genes) (Tsuji et al. 2007). Several hypotheses have been proposed to explain why more than half of tRNA genes has been lost and how to compensate for the absence of tRNA genes from *Selaginella* plastome (Wolfe et al. 1992) (Tsuji et al. 2007). The hypothesis that the lost tRNA are encoded in the nuclear genome and imported to the plastid from the cytosol, which is also known to occur for plant mitochondria, is most likely to be the explanation for the presence of “vestigial” tRNA genes found in *S. sanguinolenta* and *S. uncinata*. Furthermore, no tRNA genes were annotated in mitochondrial genome of *S. moellendorffii* (Hecht et al. 2011). Therefore, the absent tRNA genes in plastomes of *Selaginella* are most likely lost and imported from nucleus.

Except for the severe tRNA genes loss mentioned above, numerous protein-coding genes were also absent in *Selaginella* plastomes, especially in the species of the basal lineages (Figure 1). Five species had *ndh* gene loss to a different extant with all *ndh* genes lost in *S. vardei, S. indica*, and *S. lyallii*, ten lost and one (*ndhC*) intact in *S. tamariscina*, four lost, four pseudogenized and three intact in *S. sanguinolenta*, and 11 *ndh* genes intact in other six *Selaginella* species. The gene loss of *ndh* genes have been reported in land plant plastomes many times, such as, *Najas flexilis* (Peredo et al. 2013), *Epifagus*, (Haberhausen and Zetsche 1994), *Cuscuta* (Funk et al. 2007; Logacheva et al. 2011), *Neottia* (Feng et al. 2016; Haberle et al. 2008), Gnetales (Raubeson and Jansen 2005), Pinaceae (Wakasugi et al. 1994), and several *Erodium* species (Blazier et al. 2011b). the n*dh* genes, encoding plastid NAD(P)H-dehydrogenase complex was involved in cyclic electron flow (CEF) chain. Two independent pathways of CEF, *PGR*5-dependent and *NDH*-dependent pathways, have been characterized across land plants, with the former being the main contributor in CEF (Ruhlman et al. 2015). Therefore, the *ndh* genes that were lost in different lineages of land plants were not transferred into nucleus, but most possibly replaced by the nuclear-encoded *PGR*5-dependent pathway (Ruhlman et al. 2015). The shared character in *S. vardei, S. indica, S. lyallii* and *S. tamariscina* is the extremely dry habitat and thick wax-like components on the leaf surface, likely to reflect the strong sunlight. In this case, we presume that the loss of *ndh* genes in these five species could be related to the adaptation to the dry and light-intensive habitat. The plastome of *S. sanguinolenta* could be in the intermediate phase growing in the moderate dry habitat, and the loss and pseudogenization of *ndh* genes could be a relatively recent loss. In *S. bisulcata*, all *ndh* genes become putative pseudogenes because of the internal stop codons caused by frame shift mutation, whereas all *ndh* genes are functional in the plastome of its sister species *S. pennata* (Figure 1; Figure S4). The extensive RNA editing sites in plastomes of *Selaginella* were previously reported, and the RNA editing sites could recover the pseudogenes into functional (Oldenkott et al. 2014). However, evidence from transcriptome data is necessary to elucidate whether the putative pseudogenes in *S. bisulcata* is truly deprived of function or restored after RNA editing at the transcriptome level.

Another noteworthy event is the intron loss in Selaginellaceae (Figure 1). The intron loss in plastomes is common in land plants (Jansen and Ruhlman 2012). Most angiosperms and ferns have two introns in *clpP* gene, whereas only one intron remained in *clpP* gene of Equisetaceae in ferns (Karol et al. 2010). In *Selaginella*, however, two introns remained in *clpP* of *S. sanguinolenta* and *S. doederleinii* as in other lycophytes (Isoetaceae and Lycopodiaceae), only one intron, similar to the plastomes of Equisetaceae, remained in *clpP* gene of other reported *Selaginella* species, and no introns existed in *clpP* of *S. tamariscina, S. remotifolia, S. lyallii, S. indica*, and *S. vardei*, similar to the plastomes of *Pinus* species (Wakasugi et al. 1994), two *Silene* species (Erixon and Oxelman 2008), and grasses (Barkan 2004; Downie and Palmer 1992). The intron of *atpF* gene was lost in *S. vardei* and *S. indica*, while the intron of rpoC1 was absent in *S. lyallii*. Besides, the intron 2 of *ycf*3 gene was lost in *S. vardei, S. indica, S. lyallii*, and *S. remotifolia*. Observation of the intron loss in *atpF* was first uncovered in the plastome of cassava (Daniell et al. 2008) and the loss of rpoC1 intron was found to occur multiple times in angiosperms (Downie et al. 1996). The intron 2 loss of *ycf*3 genes represents the first documented case within the plastomes of land plants. In the case of intron loss, one mechanism has been proposed that involves recombination between a processed intron-less cDNA and the original intron-containing copy (Daniell et al. 2008). Under this situation, an apparent decrease of RNA editing sites in the neighboring regions of lost intron should be observed in genes losing intron. The multiple sequence alignments between intron-lost genes (*atpF, clpP, rpoC1*, and *ycf3*) and their homologues with intron from closely related species displayed apparent C to T change at the flanking regions of lost introns (Figure S5). Therefore, we propose that the intron loss of *clpP, atpF*, and *ycf*3 in Selaginellaceae can likely be explained by this mechanism.

### Evolutionary Trajectory of DR/IR in Selaginellaceae

Except the unconventional DR structure in *S. vardei* (Zhang et al., 2018) and *S. tamariscina* (Xu et al. 2018), we also found the DR structure existing in the plastomes of our newly sequenced seven species (*S. lyallii, S. remotifolia, S. sanguinolenta, S. tamariscina, S. doederleinii, S. moellendorffii, S. bisulcata*, and *S. pennata*), which showed that the DR structure in *Selaginella* plastomes is a remarkable character. However, the typical IR structure, existing in almost all land plants, still remains in the plastomes of *S. uncinata* and *S. hainanensis*. Furthermore, PCR confirmation of the plastome structure in 18 representative species from four subgenera sensu Zhou and Zhang (2015) (Table S2) showed that three subgenera in Selaginellaceae possess plastomes with DR structure, whereas in the most derived subg. *Heterostachys* the plastomes evolved into the typical IR structure again (Figure 2, Figure S3). Although the IR structure is ubiquitous in plastomes among land plants, the DR structure is indicated as the early diverged character and remained in plastomes of most lineages within Selaginellaceae (Figure 2). Given that plastomes from the other two families, Lycopodiaceae and Isoetaceae, of the lycophytes possess the typical IR structure (Guo et al. 2016; Karol et al. 2010; Wolf et al. 2005; Zhang et al. 2017), we suggest that the DR structure have occurred in the Selaginellaceae plastomes after the separation from Isoetaceae (Unpublished data). The occurrence of the direction change from IR in Isoetaceae to DR in Selaginellaceae was attributed to an inversion of ca. 50 kb fragment *trnF*-*trnN* spanning the complete IR_B_ region (Unpublished data). Both DR and IR structure exist in the most derived subg. *Heterostachys*, with the plastomes of *S. bisulcata* and *S. pennata* assembled into DR structure and the plastomes of *S. uncinata* and *S. hainanensis* assembled into IR structure. Two inversions were described between these two plastome structures, and one inversion of ca. 65 kb fragment *trnD*-*rpl20* spanning one copy of repeat region (IR_B_) recovered the IR structure in *S. uncinata* and *S. hainanensis* (Figure 2). Furthermore, each plastome of subg. *Heterostachys* is able to generate the IR/DR-coexisting structures owing the duplication of the large region *psb*K-*trn*Q (Figure 6), as is detailed discussed in the next section.

The result of divergence time estimation showed that the DR and IR structure (IR/DR-coexisting) occurred at about 246 Ma and 23 Ma, respectively (Figure 2). The well-known Permian-Triassic (P-T) extinction event occurred about 252 Ma ago, causing about 50% reduction in terrestrial plants diversity (McElwain and Punyasena 2007). At the P-T boundary, despite the dramatically changed terrestrial ecosystems, lycophytes, being one of the surviving plants, played a central role in repopulating the initial ecological landscape (Looy et al. 1999). Both mitochondrial and plastid genomes are more frequently subjected to alterations under specific environmental conditions (Maréchal and Brisson 2010). Given the evidence of the P-T extinction, the DR structure in Selaginellaceae is most possibly selected for better surviving the ecological upheaval (increasing anoxia, increasing aridity, and increased UV-B radiation). However, the potential mechanism for the recent recurrence of IR and the coexistence with DR structure in plastomes of Selaginellaceae need further exploration.

### Dynamic Structures of the Plastomes with DR/IR Structure of Selaginellaceae

Recombination-dependent process between homologous repeats is responsible for the highly dynamic structure of plant organelle genomes (plastomes and mitogenomes) (Maréchal and Brisson 2010; Ruhlman et al. 2017). Large repeats (> 1 kb) are able to mediate high frequency, reciprocal recombination intra- or intermolecularly and generally results in approximately equimolar amounts of the parental and recombinant forms (Arrieta-Montiel and Mackenzie 2011; Fauron et al. 1995). In genomes where both inverted and direct repeats are present, recombination activities in these different orientations will lead to drastically different genome organizations, containing various isomeric forms of the master chromosome and subgenomic molecules (Fauron et al. 1995).

The plastomes of *S. uncinata* and *S. hainanensis* were assembled into an IR structure, which we consider as master chromosome. However, another pair of ca. 2.7 kb inverted repeats spanning *psb*K-*trn*Q was identified in the LSC and SSC region, respectively (Figure S5). Therefore, recombination between IR generates an isomer with altered orientation of one single copy region, which, in turn, changes the inverted repeats of *psb*K-*trn*Q into direct (isomer I). Recombination between the copies of ca. 2.7 kb inverted repeat *psb*K-*trn*Q could change the orientation of IR into direct (DR) and generates isomer II. Both isomer I and isomer II could give rise to two subgenomic molecules through the recombination between direct repeats, respectively (Figure 6A).

The plastomes of *S. bisulcata* and *S. pennata* were assembled into DR structure, which was considered as master chromosome. However, a pair of ca. 1.8 kb inverted repeats spanning *psb*K-*trn*Q was identified in the LSC and SSC region, respectively (Figure S5). Following the recombination activities in *S. uncinata* and *S. hainanensis*, recombination between DR gives rise to two subgenomic molecules, and recombination between the copies of ca. 1.8 kb inverted repeat *psb*K-*trn*Q change the orientation of DR into inverted (IR), generating isomer I. Recombination between newly created IR in isomer I generates isomer II with altered orientation of one single copy region, which similarly changes the inverted repeats of *psb*K-*trn*Q into direct (isomer II). Finally, recombination between the copies of direct repeat *psb*K-*trn*Q in isomer II also gives rise to two different subgenomic molecules (Figure 6B). Thus, these frequent, reciprocal recombination activities created a dynamic complex heterogeneous population of plastomes in *S. uncinata, S. hainanensis, S. bisulcata*, and *S. pennata*.

However, as reported in *S. vardei* (Unpublished data), the plastomes with DR structure could only promote a master chromosome and two sets of subgenomic chromosomes at approximately equal stoichiometries by the recombination between two copies of DR within one plastome or between different molecules (Figure 6C). Either the master chromosome or subgenomic chromosomes could form head to tail concatemers of both circular and linear molecules together with branched structures through recombination between DR regions. The existence of subgenomes in species of DR-possessing plastomes have been confirmed using the PacBio reads in *S. tamariscina* (Xu et al. 2018), whereas the existence of the complex heterogeneous population of multipartite plastomes in *S. uncinata, S. hainanensis, S. bisulcata*, and *S. pennata* still need to be confirmed by long reads from PacBio or Nanopore sequencing. Considering the fact that most land plants possess plastomes with IR structure and only the early diverged lycophyte group Selaginellaceae share DR structure, plastomes with IR structure presumably have more advantageous characters for plants survival and adaptation. The coexistence of the dynamic heterogeneous plastome structures in the derived lineage is possibly in the intermediate stage, which also reached an equilibrium in plastome organization. However, the biological significance behind the diverse plastome structures, especially for adaptation to environments, and the role of nuclear-encoded, plastid-targeted genes, which control the recombination behaviors, are worth further exploration.

### Correlation between Plastome Rearrangements and Repeats in Selaginellaceae

Two main forms of rearrangements, inversions and DR/IR region expansion/contraction, constitute the main rearrangement events in plastomes of Selaginellaceae. Plastome organizations of basal lineages (e.g. *S. vardei, S. indica, S. lyallii, S. remotifolia*, and *S. sanguinolenta*) of Selaginellaceae showed relatively conserved gene order, whereas rearrangements mainly existed in the more evolved lineages (e.g. *S. tamariscina, S. doederleinii, S. bisulcata* and *S. uncinata*) (Figure 2; Figure S1).

The extensive rearrangement events in plastomes of Geraniaceae have shown to be correlated with high incidence of dispersed repetitive DNA (Weng et al. 2013). The correlation between number of repeats and the rearrangement distances was also detected in Selaginellaceae plastomes (BP, P=0.034, r =0.613; IVs, P=0.011, r=0.702, Figure 4B, C), with high frequency of repeats in IR/DR-coexisting plastomes (Figure 4A). However, no correlation was detected in the plastomes with DR structure (BP, P=0.498, r =-0.283; IVs, P=0.498, r =-0.283, figure not shown), suggesting that the repeats in Selaginellaceae are more associated with the structure change from DR to IR/DR coexistence. The coexistence of few repeats and DR structure in *Selaginella* plastomes, as reported in *S. vardei* and *S. indica* (Zhang et al., 2018) might have conferred an advantage to maintain the plastome stability and have been selected. On the other hand, the occurrence of repeats, especially the large one (*psb*K-*trn*Q), in the IR/DR-coexisting plastomes are responsible for the hypothesized dynamic plastome complexity, which has reached an equilibrium state in order to maintain stability.

### Correlation between Plastome Rearrangements and Nucleotide Substitution Rate in Selaginellaceae

The significantly low d*S* value of genes of IR/DR-coexisting plastomes indicated that more efficient recombination activities, functioning as gene conversion mechanism and occurring within the single copy regions, consequently decrease the d*S* value in IR/DR-coexisting plastomes (Ruhlman and Jansen 2014; Ruhlman et al. 2017). However, the genes from IR/DR-coexisting plastomes exhibited accelerated d*N* and d*N*/d*S* compared to the genes of DR-possessing plastomes suggesting that genes in species with IR/DR-coexisting plastomes is presumably subject to much higher selective pressures, which may require fixation of functionally important mutation (Bousquet et al. 1992). For species with DR-possessing plastomes, the difference of d*S* between genes inside IV and outside IV was not significant (P= 0.042), showing the 50 kb inversion causing DR structure did not have significant influence on synonymous substitution rate. The significantly low d*N* value and d*N*/d*S* of genes outside IV is possibly correlated with the genes themselves, which usually encode the main subunit of photosynthesis-necessary proteins and under strong selection pressure (Table S6).

## Acknowledgments

This work was supported by the National Natural Science Foundation of China (grant number 31670205, 31770237). The authors would like to thank Chang-Hao Li for help with data analyses, Shi-Liang Zhou, Wen-Pan Dong and Jong-Soo Kang for helpful discussions, Nawal Shrestha for helpful revision and polishing the whole manuscript.

## Supplementary Material

**Figure S1. Detailed plastid genome structures of Lycophyte species. HL**, *Huperzia lucidula*; **HS**, *H. serrata*; **HJ**, *H. javanica*; **IF**, *Isoetes flaccida*; **SV**, *Selaginella vardei*; **SI**, *S. indica*; **SL**, *S. lyallii*; **SR**, *S. remotifolia*; **SS**, *S. sanguinolenta*; **ST**, *S. tamariscina*; **SD**, *S. doederleinii*; **SM**, *S. moellendorffii*; **SP**, *S. pennata*; **SB**, *S. bisulcata*; **SU**, *S. uncinata*; **SH**, *S. hainanensis*. Genes are colored by function. Black star marks the occurrence of DR structure, grey star marks the occurrence of IR/DR-coexisting structure in Selaginellaceae. Orange arrows in DR/IR region of plastomes show the contraction and expansion.

**Figure S2 PCR gels of the representative species in subgenera. a**, species from subg. *Heterstachys*. **8203**, *S. helferi*; **7833**, *S. picta*; **2415**, *S. mairei*; **7187**, *S. delicatula*; **6548**, *S. willdenowii*. **1**, *ycf*2-*ndh*J; **2**, *rpl*2-*rpl*23; **3**, *ycf*3-*rps*4; **4**, *trn*D-*pet*N; **5**, *ndh*F-*rps*4; **6**, *rpl*2-*rps*7. **b**, species from subg. *Stachygynandrum*. **7867**, *S. commutata*; **7899**, *S. guihaia*; **7895**, *S. rolandiprincipis*; **8176**, *S. scabrifolia*; **7863**, *S. biformis*; **7643**, *S. davidii*; **8201**, *S. erythropus*; **2016015**, *S. gebauriana*; **6050**, *S. involvens*. **1**, *ycf*3-*rps*4; **2**, *rrn*16-*rpl*23; **3**, *trn*F-*chl*L; **4**, *ndh*F-*rps*4; **5**, *rrn*16-*rps*7. **c**, species from subg. *Pulviniela*. **519**, *S. pulvinata*; **7644**, *S. stauntoniana*. **1**, *ycf*3-*ycf*3; **2**, *rrn*16-*rpl*23; **3**, *ndh*C-*chl*L; **4**, *ccs*A*-ycf*3; **5**, *rrn*16-*rps*7. **d**, species from subg. *Boreoselaginella*. **179**, *S. nummularifolia*; **7833**, *S. rossii*. **1**, *rps*4-*rrn*5; **2**, *ndh*B-*rpl*2; **3**, *trn*F-*chl*L; **4**, *ndh*F-*rrn*5; **5**, *trn*L-*trn*C.

**Figure S3 Comparisons of *ndh* genes between *S. bisulcata* and *S. pennata*.**

**Figure S4 Comparisons of intron-loss genes among closely related species.**

**Figure S5. Distribution of repeats in plastomes of newly sequenced *Selaginella* species. Supplementary tables:**

Table S1. Technical details of the Illumina datasets and genome assemblies.

Table S2 PCR confirmation for plastomes structure of representative species of four genera.

Table S3. Primers newly designed for PCR amplifications of the sampled representative species of the five main lineages in Selaginellaceae.

Table S4. The permutation of number coded Locally Collinear Block (LCB) for each plastome. Negative number indicates an inversion of the given LCB.

Table S5. Locally Collinear Blocks identified by Mauve alignment of 12 plastomes of lycophytes.

Table S6. Genes inside and outside ca. 50 kb inversion of plastomes with DR structure.

