## Supplementary figures and images for "The Unique Evolutionary Trajectory and Dynamic Conformations of DR and IR/DR- coexisting Plastomes of the Early Vascular Plant Selaginellaceae (Lycophyte)"

### Figure S1

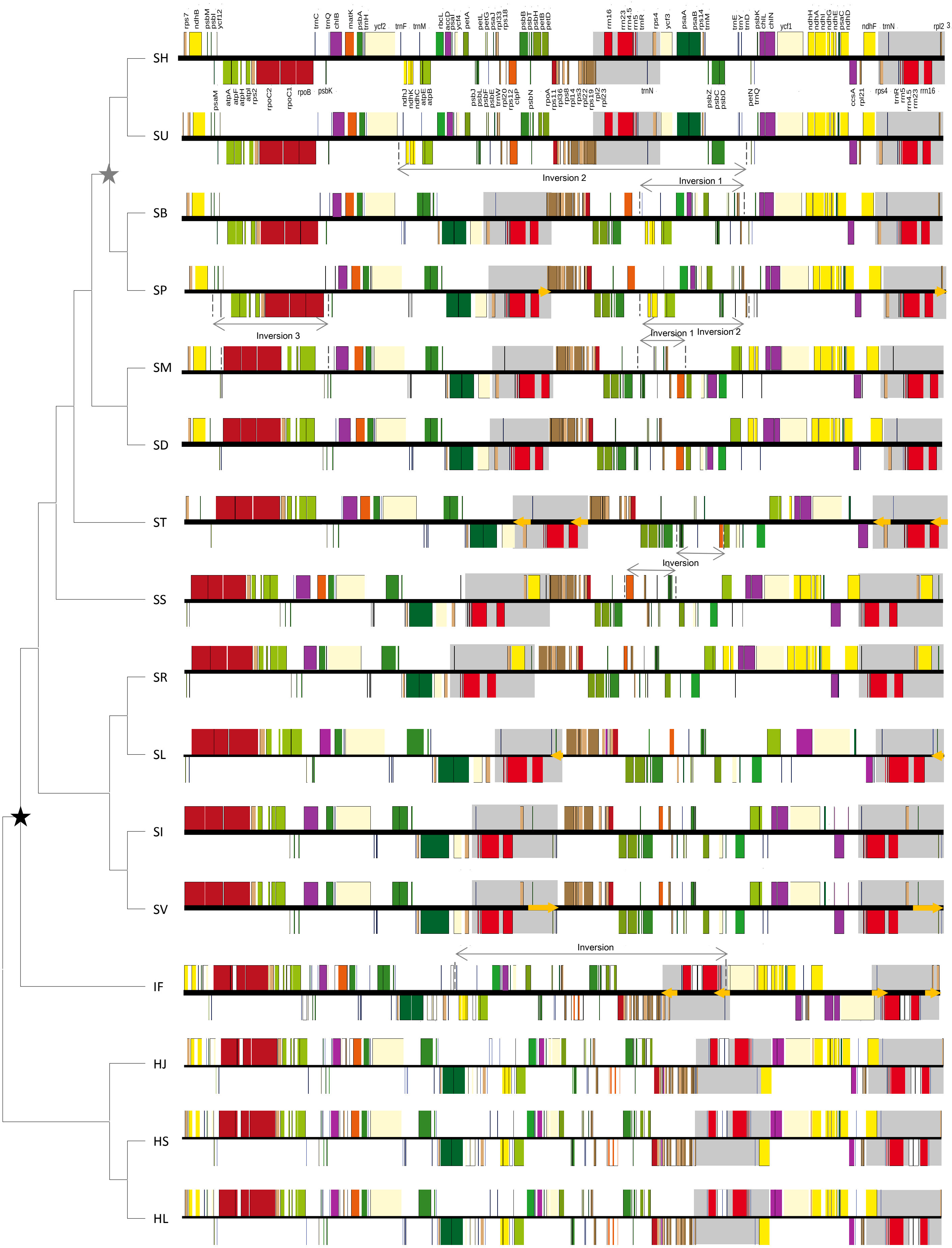

### Figure S2

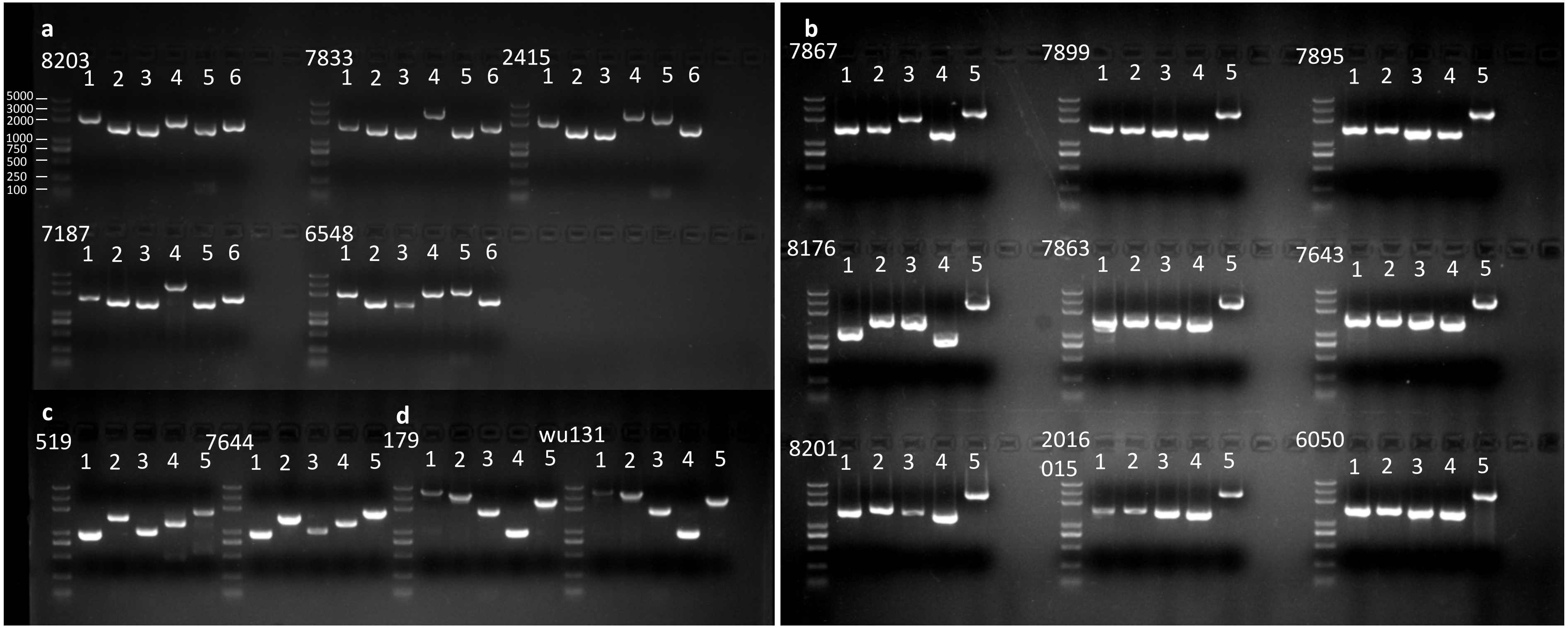

### Figure S5

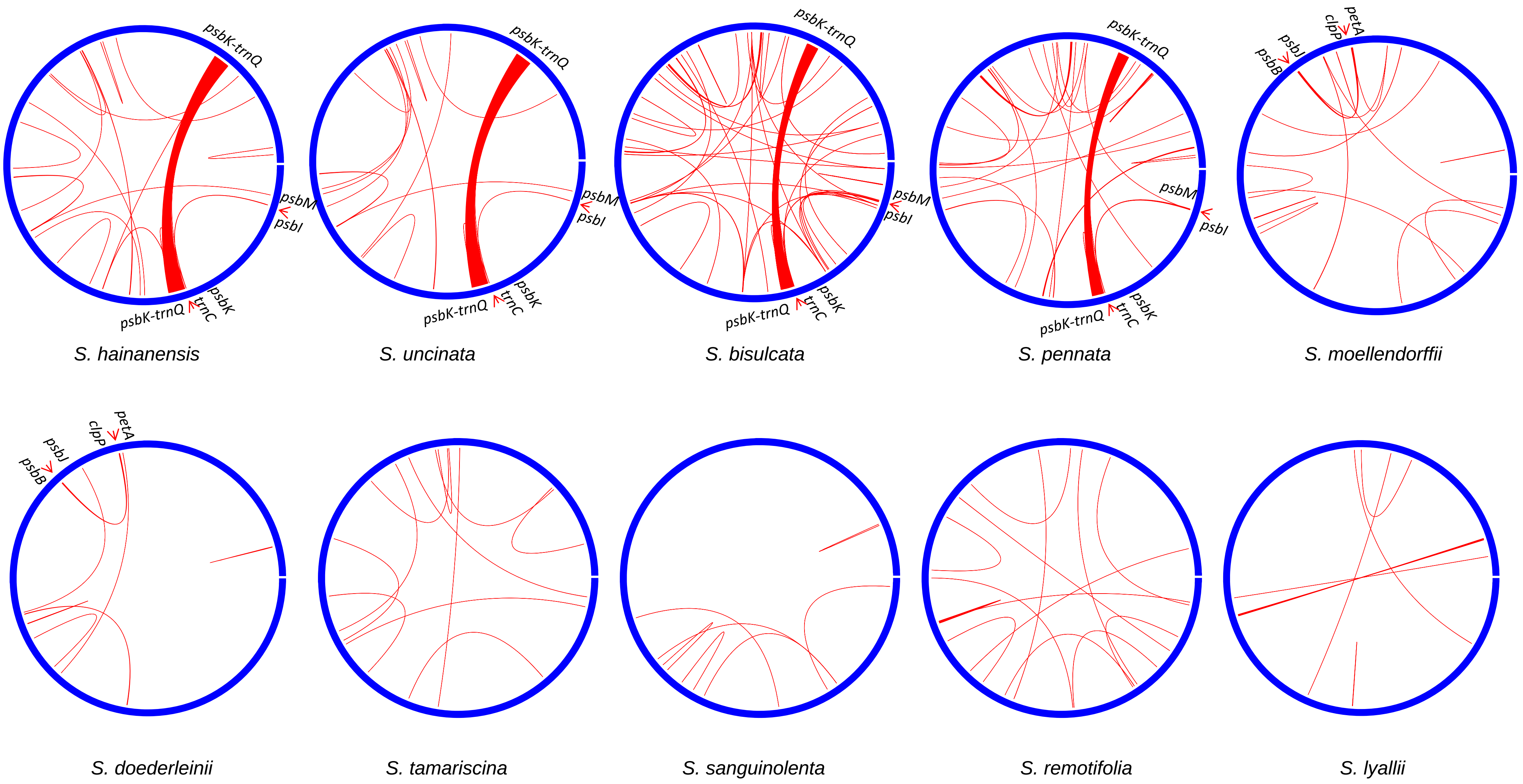
