## Supplementary material for "The Unique Evolutionary Trajectory and Dynamic Conformations of DR and IR/DR- coexisting Plastomes of the Early Vascular Plant Selaginellaceae (Lycophyte)": Figure S3

| *ndh* A(bp) | 16 | 106 | 116 | 224 | 242 | 292 | 389 | 1482 | 1487 | 1499 | 1580 | 1595 | 1610 | 1737 | 1781 | 1802 | 1888 | 1911 | 1974 | 1993 |
|---|---|---|---|---|---|---|---|---|---|---|---|---|---|---|---|---|---|---|---|---|
| *S. bisulcata* | ------ | ---- | * | 3 bp | ----- | * | 4bp | 4bp | * | * | * | * | --- | ------------ | * | * | ---- | 4bp | * | 3 bp |
| *S. pennata* | 6 bp | 4 bp | V | --- | 5 bp | S | ---- | ---- | E | K | E | D | 3 bp | 12 bp | N | D | 4 bp | ---- | N | --- |
| | exon 1 | | | | | | | exon 2 | | | | | | | | | | | | |

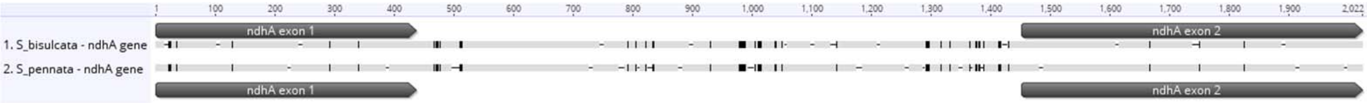


| *ndh* B (bp) | 106 | 211 | 269 | 323 | 347 | 359 | 392 | 407 | 488 | 563 | 590 | 602 | 653 | 1845 | 1879 | 2039 | 2086 | 2114 | 2151 | 2191 | 2301 |
|---|---|---|---|---|---|---|---|---|---|---|---|---|---|---|---|---|---|---|---|---|---|
| *S. bisulcata* | 3 bp | ---- | * | * | * | * | * | * | * | --- | * | * | * | ----- | * | ---- | 7 bp | ---- | 1 bp | ------ | * |
| *S. pennata* | --- | 4 bp | L | V | L | L | L | V | M | 3 bp | L | V | I | 5 bp | S | 4 bp | ------- | 4 bp | - | 6 bp | N |
| | exon 1 | | | | | | | | | | | | | exon 2 | | | | | | | |

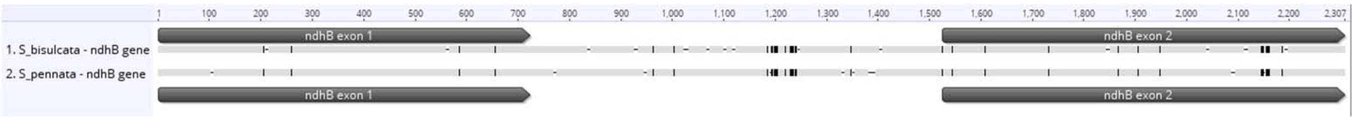


| *ndhC* (bp) | 55 | 70 | 132 | 178 | 289 |
|---|---|---|---|---|---|
| *S. bisulcata* | 2 bp | * | 2 bp | 5 bp | 4 bp |
| *S. pennata* | -- | A | -- | ----- | ---- |

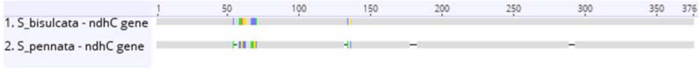


| *ndhE* (bp) | 178 | 190 | 298 |
|---|---|---|---|
| *S. bisulcata* | 2 bp | * | * |
| *S. pennata* | -- | A | L |

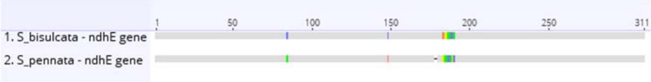


| *ndh* D(bp) | 25 | 76 | 145 | 265 | 323 | 335 | 368 | 374 | 389 | 483 | 514 | 516 | 627 | 760 | 767 | 860 | 869 | 899 | 923 | 980 | 989 | 998 | 1006 | 1088 | 1118 | 1176 | 1257 | 1362 | 1380 |
|---|---|---|---|---|---|---|---|---|---|---|---|---|---|---|---|---|---|---|---|---|---|---|---|---|---|---|---|---|---|
| *S. bisulcata* | 3 bp | 3 bp | --- | ---- | * | --------- | * | * | ---- | --- | 3 bp | * | ---- | ---- | * | * | * | * | * | * | * | * | 5 bp | * | ---- | --- | ------ | 3 bp | 3 bp |
| *S. pennata* | --- | --- | 3 bp | 4 bp | L | 9 bp | L | L | 4 bp | 3 bp | --- | K | 4 bp | 4 bp | L | L | V | L | V | I | T | I | ----- | R | 4 bp | 3 bp | 6 bp | --- | --- |

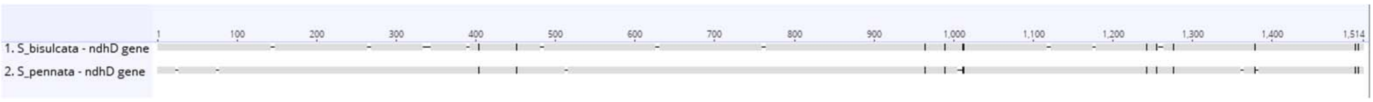


| *ndh* F (bp) | 259 | 319 | 331 | 409 | 434 | 513 | 518 | 626 | 650 | 656 | 699 | 714 | 880 | 883 | 955 | 967 | 1006 | 1152 | 1174 | 1243 | 1296 | 1334 | 1389 | 1467 | 1522 | 1576 | 1669 | 1696 | 1707 |
|---|---|---|---|---|---|---|---|---|---|---|---|---|---|---|---|---|---|---|---|---|---|---|---|---|---|---|---|---|---|
| *S. bisulcata* | 3 bp | 5 bp | * | ---- | * | 3 bp | * | * | * | ---- | --- | ------- | * | * | * | * | * | 3 bp | * | -------- | ----- | ---- | 6 bp | ---- | * | --- | * | * | 3 bp |
| *S. pennata* | --- | ----- | M | 4 bp | N | --- | R | E | E | 4 bp | 3 bp | 7 bp | M | M | L | L | L | --- | L | 8 bp | 5 bp | 4 bp | ------ | 4 bp | L | 3 bp | M | I | --- |

| | 1759 | 1771 | 1797 | 1924 | 1966 | 1982 | 1994 | 2071 | 2074 | 2240 |
|---|---|---|---|---|---|---|---|---|---|---|
| *S. bisulcata* | * | * | 4 bp | 4 bp | * | 3 bp | 5 bp | 4 bp | *N | 4 bp |
| *S. pennata* | L | L | ---- | ---- | D | --- | ----- | ---- |  | ---- |

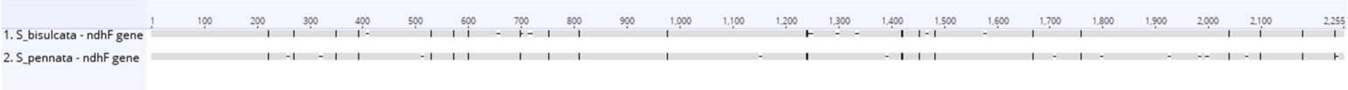


| *ndhG* (bp) | 34 | 67 | 98 | 161 | 203 | 239 | 263 | 412 | 476 | 508 | 534 |
|---|---|---|---|---|---|---|---|---|---|---|---|
| *S. bisulcata* | 3 bp | ---- | * | * | ------ | --- | ----- | ------- | ---- | 3 bp | * |
| *S. pennata* | --- | 4 bp | T | L | 6 bp | 3 bp | 5 bp | 7 bp | 4 bp | --- | E |

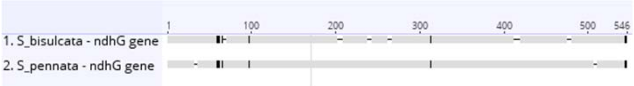


| *ndhI* (bp) | 178 | 196 | 212 | 238 | 262 | 284 | 493 | 532 |
|---|---|---|---|---|---|---|---|---|
| *S. bisulcata* | 4 bp | * | ------ | * | --- | 5 bp | 8 bp | * |
| *S. pennata* | ---- | E | 6 bp | D | 3 bp | ----- | -------- | V |

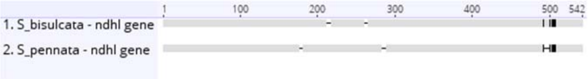


| *ndh* H (bp) | 52 | 97 | 115 | 160 | 259 | 410 | 431 | 464 | 506 | 515 | 520 | 645 | 678 | 807 | 846 | 894 | 909 | 944 | 1128 | 1147 | 1165 |
|---|---|---|---|---|---|---|---|---|---|---|---|---|---|---|---|---|---|---|---|---|---|
| *S. bisulcata* | 1 bp | * | * | * | ---- | * | 4 bp | * | 1 bp | * | 4 bp | ---- | --------- | * | * | * | --- | 4 bp | ---- | * | * |
| *S. pennata* | - | E | E | E | 4 bp | E | ---- | N | - | V | ---- | 4 bp | 9 bp | M | L | M | 3 bp | ---- | 4 bp | M | I |

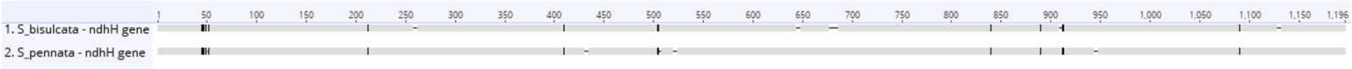


| *ndhJ* (bp) | 42 | 68 | 84 | 111 | 147 | 192 | 201 | 305 | 316 | 364 | 437 | 464 |
|---|---|---|---|---|---|---|---|---|---|---|---|---|
| *S. bisulcata* | 3 bp | -------- | * | * | * | * | ----- | * | 3 bp | 3 bp | * | -- |
| *S. pennata* | --- | 8 bp | E | D | E | N | 5 bp | I | --- | --- | L | 2 bp |

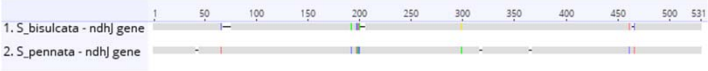


| *ndhK* (bp) | 196 | 235 | 328 | 335 | 338 | 464 | 521 | 616 |
|---|---|---|---|---|---|---|---|---|
| *S. bisulcata* | --- | ------ | ---- | * | * | * | * | 5 bp |
| *S. pennata* | 3 bp | 6 bp | 4 bp | L | V | V | I | ----- |

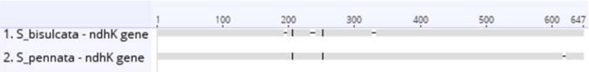
