## Supplementary material for "The Unique Evolutionary Trajectory and Dynamic Conformations of DR and IR/DR- coexisting Plastomes of the Early Vascular Plant Selaginellaceae (Lycophyte)": Figure S4

| *rpo* C1 (bp) | ... | 257 | 258 | 259 | 275 | 282 | 284 | 291 | 296 | 318 | 347 | 387 | 419 | 432 | ... | 1242 | 1245 | 1258 | 1276 | 1309 | 1318 | 1330 | 1338 | 1339 | 1341 | 1351 |
|---|---|---|---|---|---|---|---|---|---|---|---|---|---|---|---|---|---|---|---|---|---|---|---|---|---|---|
| *S. vardei* | ... | C | C | C | C | C | C | C | C | C | C | C | C | C | ... | C | C | C | C | C | C | C | C | C | C | C |
| *S. indica* | ... | C | C | C | C | C | C | C | C | C | C | C | C | C | ... | C | C | C | C | C | C | C | C | C | C | C |
| *S. remotifolia* | ... | T | C | T | T | C | T | C | C | C | T | C | C | C | ... | C | T | C | T | T | C | C | T | T | C | C |
| *S. lyallii* | ... | T | T | T | T | T | T | T | T | T | T | T | T | T | ... | T | T | T | T | T | T | T | T | T | T | T |

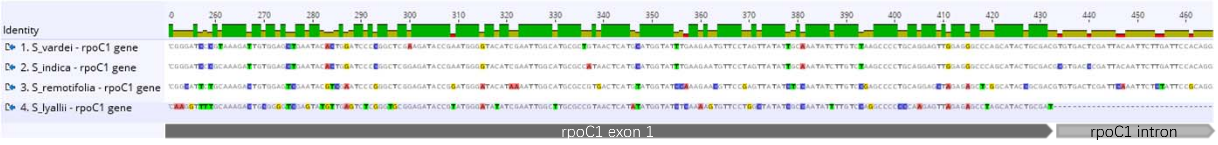


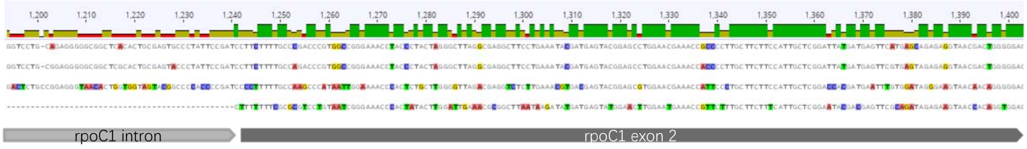


| *atp* F (bp) | ... | 11 | 30 | 42 | 51 | ... | 905 | 916 | 972 | 1024 | 1026 | 1027 |
|---|---|---|---|---|---|---|---|---|---|---|---|---|
| *S. vardei* | ... | T | T | T | T | ... | T | T | T | T | T | T |
| *S. indica* | ... | T | T | T | T | ... | T | T | T | T | T | T |
| *S. remotifolia* | ... | C | C | C | C | ... | C | C | C | C | C | C |
| *S. lyallii* | ... | C | C | C | C | ... | C | C | C | C | C | C |

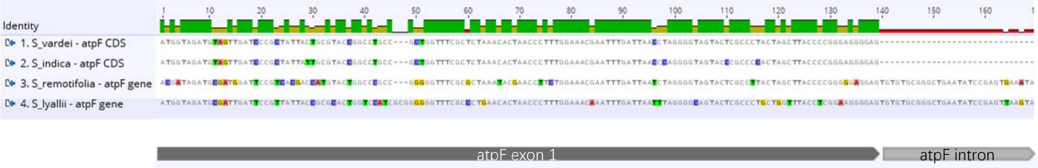


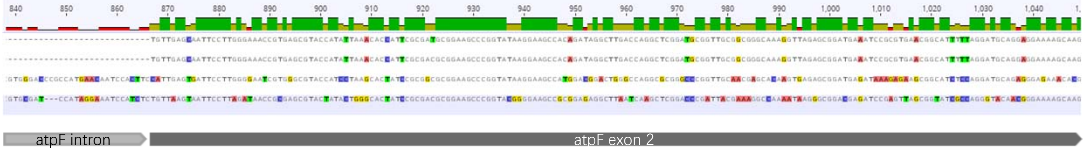


| *ycf* 3 (bp) | ... | 1033 | 1067 | 1075 | 1088 | 1094 | 1108 | 1120 | 1129 | 1183 | 1186 | 1217 | 1230 | 1255 | ... | 2222 | 2244 | 2257 | 2274 | 2314 | 2328 |
|---|---|---|---|---|---|---|---|---|---|---|---|---|---|---|---|---|---|---|---|---|---|
| *S. vardei* | ... | T | C | T | C | T | T | T | T | T | T | T | T | T | ... | T | T | T | T | T | T |
| *S. indica* | ... | T | T | T | T | T | T | T | T | T | T | T | T | T | ... | T | T | T | T | T | T |
| *S. remotifolia* | ... | T | T | T | T | T | T | T | T | T | T | T | T | T | ... | T | T | T | T | T | T |
| *S. lyallii* | ... | T | T | T | T | T | T | T | T | T | T | T | T | T | ... | T | T | T | T | T | T |
| *S. sanguinolenta* | ... | C | C | C | C | C | C | T | C | C | C | T | C | C | ... | T | C | C | C | T | C |
| *S. tamariscina* | ... | T | C | C | C | C | T | T | T | C | T | T | C | T | ... | T | T | T | T | T | C |
| *S. doederleinii* | ... | C | T | C | C | T | C | C | C | C | C | C | C | C | ... | C | C | C | C | C | C |
| *S. moellendorffii* | ... | C | T | C | C | T | C | C | C | C | C | C | C | C | ... | C | C | C | C | C | C |

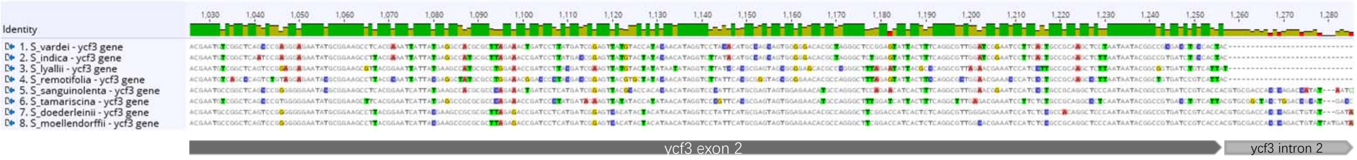


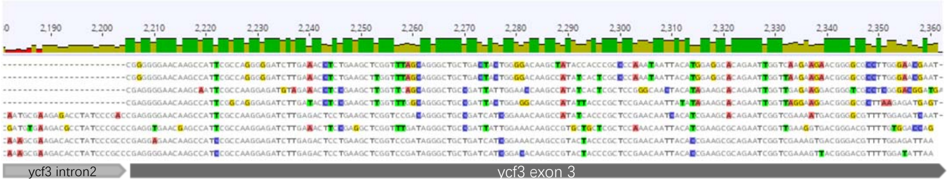

| *clp* P (bp) | 58 | 68 | 70 | … | 1006 | 1042 | 1044 | 1048 | 1059 | 1078 | 1137 | 1168 | 1119 | 1193 | … | 1951 | 1960 | 2026 | 2113 | 2131 | 2136 |
|---|---|---|---|---|---|---|---|---|---|---|---|---|---|---|---|---|---|---|---|---|---|
| *S. vardei* | T | T | T | … | T | T | T | T | T | T | T | T | T | C | … | C | C | C | T | C | T |
| *S. indica* | T | T | T | … | T | T | T | T | T | T | T | T | T | C | … | C | C | C | T | C | T |
| *S. remotifolia* | T | T | T | … | T | T | T | T | T | T | T | T | T | T | … | T | C | C | T | C | T |
| *S. lyallii* | T | T | T | … | T | T | T | T | T | T | T | T | T | C | … | T | C | A | A | T | T |
| *S. tamariscina* | T | C | T | … | T | T | T | T | T | T | T | T | T | C | … | T | C | C | T | T | T |
| *S. moellendorffii* | C | C | C | … | T | T | T | T | T | T | T | T | C | C | … | C | C | A | C | C | C |
| *S. bisulcata* | C | C | C | … | C | C | C | C | C | C | C | C | C | T | … | C | T | T | C | T | C |
| *S. pennata* | C | C | C | … | C | C | C | C | C | C | C | C | C | T | … | C | T | T | C | T | C |
| *S. hainanensis* | C | C | C | … | C | C | C | C | C | C | C | C | C | T | … | C | T | T | C | T | C |
| *S. uncinata* | C | C | C | … | C | C | C | C | C | C | C | C | C | T | … | C | T | T | C | T | C |
| *S. doederleinii* | C | C | C | … | C | C | C | C | C | C | C | C | C | C | … | C | C | C | C | C | C |
| *S. sanguinolenta* | C | C | C | … | C | C | C | C | C | C | C | C | C | C | … | C | C | C | C | C | C |

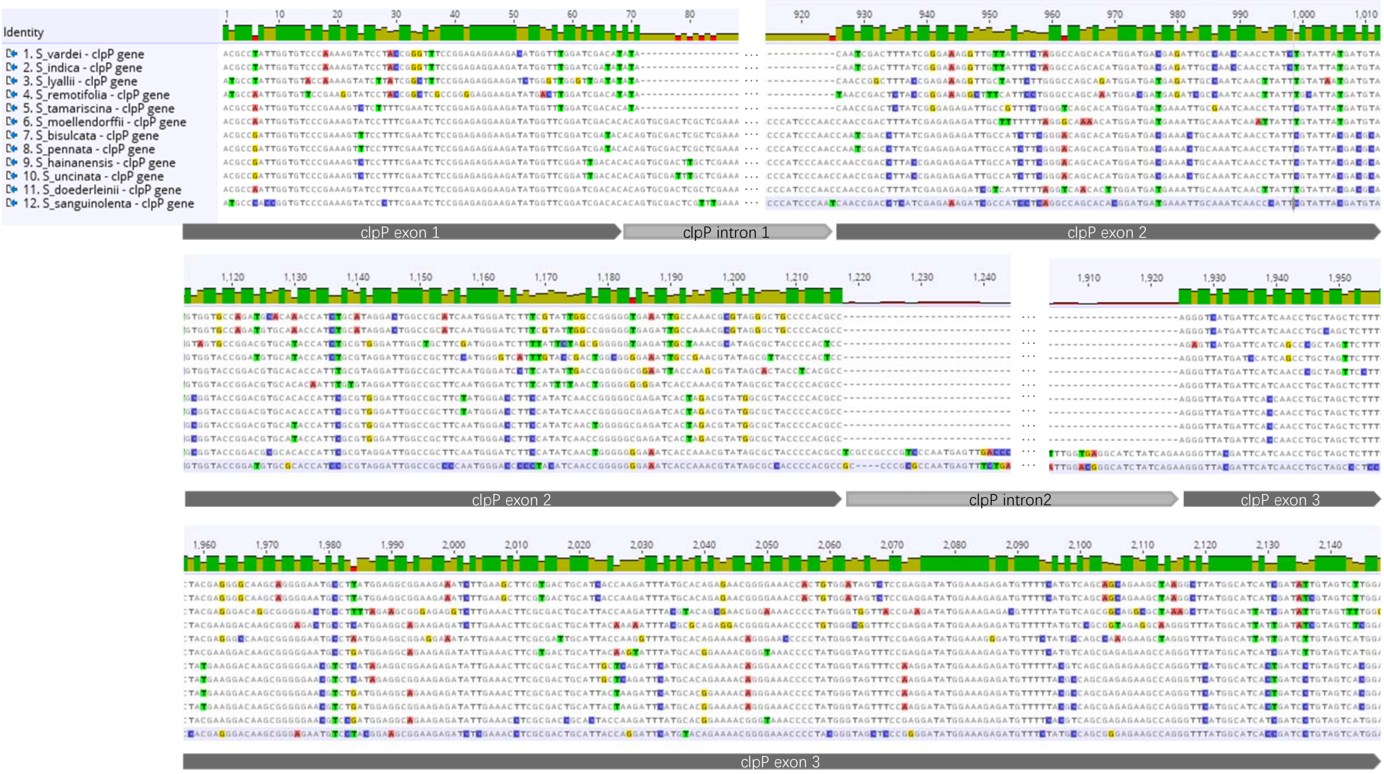
