## Supplemental Data 1 for "The Unique Evolutionary Trajectory and Dynamic Conformations of DR and IR/DR- coexisting Plastomes of the Early Vascular Plant Selaginellaceae (Lycophyte)"

| **Species** | **Genbank No.** | **Voucher** | **Clean Reads** | **No. Reads of plastome** | **Percentage** | **Coverage** |
| --- | --- | --- | --- | --- | --- | --- |
| *S. uncinata* | MG272483 | Zhang H. R. et al. 7379 (PE) | 18,426,580 | 454,363 | 2.50% | 677 |
| *S. hainanensis* | MH598533 | Zhang X. C. 7746 (PE) | 17,045,586 | 1,693,110 | 9.90% | 2224 |
| *S. bisulcata* | MH598531 | Zhang X. C. 8008 (PE) | 11,722,664 | 488,798 | 4.20% | 688 |
| *S. pennata* | MH598534 | Jin X. H. PT5014 (PE) | 36,713,270 | 5,645,918 | 15.40% | 8672 |
| *S. moellendorffii* | MG272484 | Zhang H. R. et al. 7332 (PE) | 40,787,170 | 2,010,545 | 4.90% | 3039 |
| *S. doederleinii* | MH598532 | Zhang X. C. 7797 (PE) | 34,063,566 | 605,714 | 1.80% | 874 |
| *S. tamariscina* | MH598537 | Liu B. D. sn. (PE) | 18,522,784 | 1,147,152 | 6.20% | 1837 |
| *S. sanguinolenta* | MH598536 | Zhang Z. S. 434 (PE) | 14,827,912 | 2,250,434 | 15.20% | 2812 |
| *S. remotifolia* | MH598535 | Zhang X. C. et al. 7244 (PE) | 14,446,590 | 1,025,598 | 7.10% | 1449 |
| *S. lyallii* | MK156800 | Zhang X. C. 9143 (PE) | 16,434,706 | 1,001,700 | 6.11% | 1360 |

Table S1 Technical details of the Illumina datasets and genome assemblies.

Table S2 PCR confirmation for plastomes structure of representative species of five subgenera.

|  | **Species** | **Collect No.** | **Location** | **Plastome** |
| --- | --- | --- | --- | --- |
| **subg. *Heterostachys*** | | | |  |
|  | *S. delicatula* | 7187 | Guizhou, China | with IR |
|  | *S. helferi* | 8203 | Guangxi, China | with IR |
|  | *S. mairei* | 2415 | Sichuan, China | with IR |
|  | *S. picta* | 7833 | Guizhou, China | with IR |
|  | *S. willdenowii* | 6548 | Singapore | with IR |
| **subg. *Stachygynandrum*** | | | |  |
|  | *S. commutata* | 7867 | Guangxi, China | with DR |
|  | *S. guihaia* | 7899 | Guangxi, China | with DR |
|  | *S. rolandiprincipis* | 7895 | Guangxi, China | with DR |
|  | *S. scabrifolia* | 8176 | Hainan, China | with DR |
|  | *S. biformis* | 7863 | Guangxi, China | with DR |
|  | *S. davidii* | 7643 | Beijing, China | with DR |
|  | *S. erythropus* | 8201 | Guangxi, China | with DR |
|  | *S. gebauriana* | 2016015 | Guizhou, China | with DR |
|  | *S. involvens* | 6050 | Taiwan, China | with DR |
| **subg. *Pulviniela*** | | | |  |
|  | *S. pulvinata* | 519 | Xizang, China | with DR |
|  | *S. stauntoniana* | 7644 | Beijing, China | with DR |
| **subg. *Boreoselaginella*** | | | |  |
|  | *S. nummularifolia* | 179 | Xizang, China | with DR |
|  | *S. rossii* | wu131 | Jilin, China | with DR |

Table S3 Primers newly designed for PCR amplifications of the representative species from the five main lineages of Selaginellaceae.

| **Primers** | **Sequences (5'to3')** |
| --- | --- |
| u1-ycf2-F | TTGTTCGCTCTGAATTGC |
| u1-*ndhJ*-R | GACCTTTCATGAAAGTCATCCACG |
| u2-*rpl2*-F | GATGTAGCCGATTGACCTT |
| u2-*rpl23*-R | AGCCGAATCCGAGTAAGA |
| u3-*ycf3*-F | CGAACTCACCTTCACCAA |
| u3-*rps4*-R | CGGATGCTATTGCTGCTA |
| u4-*trnD*-F | GTAGTTCAATCGGTGAGAGT |
| u4-*petN*-R | GACGGTAGTTCTTACATCTTC |
| u5-*ndhF*-F | GACCTGTTGCTGAATTACTC |
| u5-*rps4*-R | TAGCCGTAGAATATCATCCTC |
| u6-*rpl2*-F | TCGTAGTGCGTACATAAGTC |
| u6-*rps7*-R | ATGCGGAACGGATAATGG |
| d1-*ycf3*-F | CGCTACAATGGTAAGAGTCT |
| d1-*rps4*-R | TCGGTTAGGTATGGCTTCT |
| d2-*rrn16*-F | TCCTCCTTCTGCTCATCA |
| d2-*rpl23*-R | GCCTTGTCCGTAGATACC |
| d3-*trnF*-F | TCAATCCTCCGCTCTACC |
| d3-*chlL*-R | GACCAACTCCTCCGTAAC |
| d4-*ndhF*-F | CGTCAATGGCGTTGGTAT |
| d4-*rps4*-R | GTAGAATATCATCCTCGTCAAG |
| d5-*rrn16*-F | TCCTCCTTCTGCTCATCA |
| d5-*rps7*-R | CACTCTACGGATTGCTTGA |
| t1-*ycf3*-F | GGCATGTTCTCCACTACTC |
| t1-*ycf3*-R | AGCATCCACCTATCACAGA |
| t2-*rrn16*-F | GACTGATACGATGATGACTTC |
| t2-*rpl23*-R | GCTCGGCATTGGAACTATA |
| t3-*ndhC*-F | TTGGAATGGTCGGTCTAATC |
| t3-*chlL*-R | TTACGACTTGTGGTTGGTT |
| t4-*ccsA*-F | TGAAGAGCGGTGACTAGA |
| t4-*ycf3*-R | GCTTGGTGAAGGTGAGTT |
| t5-*rrn16*-F | GACTGATACGATGATGACTTC |
| t5-*rps7*-R | TGACGAGGAACGGATAATG |
| s1-*rps4*-F | GGAGTGGATTGGTTCGTAT |
| s1-*rrn5*-R | TGGTGCCTTAGTCGTAGA |
| s2-*ndhB*-F | AGCCACCAGAATCTTCAATA |
| s2-*rpl2*-R | GTCAGAATAGCATCTCCAATC |
| s3-*trnF*-F | TCAATCCTCCGCTCTACC |
| s3-*chlL*-R | TATTACGACTTGTGGTTGGT |
| s4-*ndhF*-F | ATGCGATGAACGGATATACA |
| s4-*rrn5*-R | ACTCTACTGCGGTTACGA |
| s5-*trnL*-F | CTTGGTGGTGGAATGGTAG |
| s5-*trnC*-R | CGGTAAGGCAGAGGATTG |

Table S4. The permutation of number coded Locally Collinear Block (LCB) for each plastome. Negative number indicates an inversion

| *H_serrata* | 1 | 2 | 3 | 4 | 5 | 6 | 7 | 8 | 9 | 10 | 11 | 12 | 13 | 14 | 15 | 16 | 17 | 18 | 19 |  |
| --- | --- | --- | --- | --- | --- | --- | --- | --- | --- | --- | --- | --- | --- | --- | --- | --- | --- | --- | --- | --- |
| *I_flaccida* | 2 | 3 | 4 | 5 | 6 | 7 | 9 | 10 | 11 | 12 | 13 | 14 | 15 | 16 | -1 | 17 | 19 | -18 | -8 |  |
| *S_vardei* | 4 | 5 | 6 | 7 | 8 | 9 | 10 | 11 | -17 | 1 | 2 | 3 | -16 | -15 | -14 | -13 | -12 | 18 | 19 |  |
| *S_indica* | 4 | 5 | 6 | 7 | 8 | 9 | 10 | 11 | -17 | 1 | 2 | 3 | -16 | -15 | -14 | -13 | -12 | 18 | 19 |  |
| *S. remotifolia* | 3 | 4 | 5 | 6 | 7 | 8 | 9 | 10 | 11 | -17 | 1 | 2 | -16 | -15 | -14 | -13 | -12 | 18 | 19 |  |
| *S. lyallii* | 3 | 4 | 5 | 6 | 7 | 8 | 9 | 10 | 11 | -17 | 1 | 2 | -16 | -15 | -14 | -13 | -12 | 18 | 19 |  |
| *S. sanguinolenta* | 3 | 4 | 5 | 6 | 7 | 8 | 9 | 10 | 11 | -17 | 1 | 2 | -16 | -15 | -14 | -13 | -12 | 18 | 19 |  |
| *S. tamariscina* | 1 | 2 | 3 | 4 | 5 | 6 | 7 | 8 | 9 | 10 | 11 | -17 | -16 | 14 | 15 | -13 | -12 | 18 | 19 |  |
| *S. doederleinii* | 1 | 2 | 3 | 4 | 5 | 6 | 7 | 8 | 9 | 10 | 11 | -17 | -16 | 14 | 15 | -13 | -12 | 18 | 19 |  |
| *S. moellendorffii* | 1 | 2 | 3 | 4 | 5 | 6 | 7 | 8 | 9 | 10 | 11 | -17 | -16 | 14 | 15 | -13 | -12 | 18 | 19 |  |
| (Intermediate) | 1 | 2 | 3 | -5 | -4 | 6 | 7 | 8 | 9 | 10 | 11 | -17 | -16 | -15 | -14 | -13 | -12 | 18 | 19 |  |
| *S. pennata* | 1 | 2 | -5 | -4 | 6 | 7 | 8 | 9 | 10 | 11 | -17 | -16 | -15 | 12 | 13 | 14 | 3 | -6 | 18 | 19 |
| *S. bisulcata* | 1 | 2 | -5 | -4 | 6 | 7 | 8 | 9 | 10 | 11 | -17 | -16 | -15 | 12 | 13 | 14 | 3 | -6 | 18 | 19 |
| (Intermediate) | 1 | 2 | -5 | -4 | 6 | 7 | 8 | 9 | 10 | 11 | -17 | -16 | -15 | -14 | -13 | -12 |  |  |  |  |
| *S. uncinata* | 1 | 2 | -5 | -4 | 6 | 7 | 8 | 12 | 13 | 14 | 15 | 16 | 17 | -11 | -10 | -9 | 3 | -6 | 18 | 19 |
| *S. hainanensis* | 1 | 2 | -5 | -4 | 6 | 7 | 8 | 12 | 13 | 14 | 15 | 16 | 17 | -11 | -10 | -9 | 3 | -6 | 18 | 19 |

of the given LCB

Table S5 Locally Collinear Blocks identified by Mauve alignment of 12 plastomes.

| LCBs | genes |
| --- | --- |
| 1 | *rps7* |
| 2 | *ndhB, trnL-CAA, psbM, ycf66* |
| 3 | *petN* |
| 4 | *trnC-GCA, rpoB, rpoC1, rpoC2, rps2, atpI, atpH, atpF, atpA, trnR-UCU* |
| 5 | *ycf12, psaM, trnS-GCU, psbI* |
| 6 | *psbK, trnQ-UUG* |
| 7 | *chlB, rps16, trnK-UUU, matK, psbA, trnH-GUG* |
| 8 | *ycf2* |
| 9 | *trnD-GUC, trnY-GUA, trnE-UUC* |
| 10 | *psbD, psbC, trnS-UGA, psbZ, trnG-UCC, trnfM-CAU, rps14, psaB, psaA, ycf3* |
| 11 | *trnS-GGA, rps4, trnT-UGU, trnL-UAA* |
| 12 | *trnF, ndhJ, ndhK, ndhC, trnV-UAC, trnM-CAU, atpE, atpB, rbcL, trnR-CCG, accD, psaI, ycf4, ycf10* |
| 13 | *petA* |
| 14 | *psbJ, psbL, psbF, psbE, petL, petG, trnW-CCA, trnP-UGG, psaJ, rpl33, rps18, rpl20* |
| 15 | *clpP* |
| 16 | *psbB, psbT, psbN, psbH, petB, petD, rpoA, rps11, rpl36, infA, rps8, rpl14, rpl16, rps3, rpl22, rps19, rpl2* |
| 17 | *trnV-GAC, rrn16, trnI-GAU, trnA-UGC, rrn23, rrn4.5, rrn5, trnR-ACG, trnN-GUU* |
| 18 | *chlL, chlN* |
| 19 | *ycf1, rps15, ndhH, ndhA, ndhI, ndhG, ndhE, psaC, ndhD, ccsA, trnL-UAG, trnP-GGG, rpl32, rpl21* |

Table S6 Genes inside and outside ca. 50 kb inversion of plastomes with DR structure.

|  | **Inside IV** | **Outside IV** |
| --- | --- | --- |
| **Genes** | *atp*B, *atp*E, *clp*P, *pet*A, *pet*B, *pet*D, *pet*G, *pet*L, *psa*I, *psa*J, *psb*B, *psb*E, *psb*F, *psb*H, *psb*J, *psb*L, *psb*N, *psb*T, *rbc*L, *rpl*16, *rpl*2, *rpl*22, *rpo*A, *rps*11, *rps*18, *rps*19, *rps*3, *rps*8 | *atp*A, *atp*F, *atp*H, *atp*I, *ccs*A, *psa*A, *psa*B, *psa*C, *psb*A, *psb*C, *psb*D, *psb*I, *psb*K, *psb*Z, *rpo*B, *rpo*C1, *ycf*12, *ycf*3 |
